# PIP4K2B is a mechanosensor and induces heterochromatin-driven nuclear softening through UHRF1

**DOI:** 10.1101/2022.03.25.485814

**Authors:** Alessandro Poli, Fabrizio A. Pennacchio, Paulina Nastaly, Andrea Ghisleni, Michele Crestani, Francesca M. Pramotton, Fabio Iannelli, Galina Beznusenko, Alexander A. Mironov, Valeria Panzetta, Sabato Fusco, Bhavwanti Sheth, Paolo A. Netti, Dimos Poulikakos, Aldo Ferrari, Nils Gauthier, Nullin Divecha, Paolo Maiuri

## Abstract

Phosphatidylinositol-5-phosphate (PtdIns5P)-4-kinases (PIP4Ks) are stress-regulated phosphoinositide kinases able to phosphorylate PtdIns5P to PtdIns(4, 5)P2. In cancer patients their expression is typically associated with bad prognosis. Among the three PIP4K isoforms expressed in mammalian cells, PIP4K2B is the one with more prominent nuclear localization. Here, we unveil the role for PIP4K2B as mechanosensor. PIP4K2B protein level, indeed, strongly decreases in cells growing on soft substrates. Its direct silencing or pharmacological inhibition, mimicking cell response to soft, triggers a concomitant reduction of the epigenetic regulator UHRF1 and induces changes in nuclear polarity, nuclear envelope tension and chromatin compaction. This substantial rewiring of the nucleus mechanical state drives YAP cytoplasmic retention and impairment of its activity as transcriptional regulator, finally leading to defects in cell spreading and motility. Since YAP signalling is essential for initiation and growth of human malignancies, our data suggest that potential therapeutic approaches targeting PIP4K2B could be beneficial in the control of the altered mechanical properties of cancer cells.

## Introduction

Nuclear mechanosensing impacts on a variety of processes, regulating genome integrity, gene expression and cell migration (1, 2). Deregulated nuclear mechanotransduction, due to nuclear structural defects like altered chromatin organization or impairment of nuclear envelope (NE) connection with the cytoskeleton, is often associated with the insurgence of different pathologies, including cancer (2–5).

Polyphosphoinositides (PPIns/PI) are lipid second messengers involved in many cellular functions, including proliferation, adhesion, cytoskeletal organisation and regulation of transcription (6). The presence of several PI-modulating enzymes in different subcellular compartments drives compartment-specific PI profiles that contribute to organelle identity and function. PI interconversion occurs also in cell nuclei, both at the NE and within the nucleus (7–9). Nuclear PIs impact on transcriptional control through regulating DNA and chromatin modifications and RNA splicing through their interaction with proteins containing PI-interacting domains (7–9). Whether they couple mechanosensing and the control of gene expression is not known. Phosphatidyl Inositol 5-Phosphate 4 Kinases (PI5P4K/PIP4K) are lipid kinases able to phosphorylate PtdIns5P to generate small pools of PtdIns(4, 5)P2 compartmentalized at specific cell organelles (10, 11). Three isoforms of PIP4K exist in mammalian cells, PIP4K2A, 2B and 2C, mainly localizing in the cytoplasm, nucleus and endomembrane compartments respectively (12–15). Deregulations of these lipid phosphotransferases have been reported in different human cancers (11, 16, 17). Among the three isoforms, PIP4K2B controls and removes PtdIns5P in the nuclear compartment, impacting on the expression of DNA-damage related genes, chromatin remodeling and protein-histone binding (7–9).

Here we show that PIP4K2B is also involved in mechanosensing, and that direct depletion of this lipid kinase, impacting on UHRF1 expression, drives both: decrease in nuclear envelope tension and chromatin decompaction. This finally induces YAP cytoplasmic retention rewiring of cytoskeletal organization and impairment of cell motility.

## Results

### PIP4K2B protein levels positively correlate with substrates stiffness

Cytoplasmic and plasma membrane PIs and PI related enzymes play both direct and indirect roles in cell mechanics (18, 19). However, no data are available about possible correlations between PIP4K signalling and cell mechanosensing. To this end, we investigated if PIP4K expression could be affected by processes involved in cell responses to substrates of different stiffness. We plated hTERT_RPE1 cells onto fibronectin (FN)-coated acrylamide gels (2.3, 30 and 55 kPa) or onto FN- or poly D-lysine (PDL)-coated glass coverslips (Figure 1A). 24 hours after cell seeding, we lysed them and assessed the expression of PIP4K2A, 2B and 2C using western blotting. While PIP4K2A and 2C were unaffected, the expression of PIP4K2B was strongly decreased in cells seeded onto soft gels (2.3kPa) and on PDL-coated coverslips (Figure 2A). Drop of PIP4K2B was confirmed in other two cell lines, MEF and Hela (Supplementary Figure 1A and 1B). Cells growing on FN coated stiff surfaces (glass or gels of 30/55 kPa) form bigger and more numerous focal adhesions and ticker actin stress fibers respect to what they do on PDL coated glass or FN coated soft surfaces (gel 2.3kPa) (20). Consistent with decrease of FA and actin stress fibers, cells seeded on soft gels or PDL-coated glass are usually characterized by a strong nuclear exclusion of the downstream nuclear effector of the Hippo signaling pathway and master regulator of mechanotransduction, Yes-associated protein 1 (YAP) (21). This event drives changes in the expression of many genes that are target of YAP nuclear activity as co-transcription factor (20). RT-qPCR analysis indicated that the decreased expression of PIP4K2B was not due to impaired RNA levels (Supplementary Figure 1C), so that the gene encoding PIP4K2B was not part of YAP target genes. On the other hand, we found that an over/night treatment of cells seeded on soft substrates (Gel 2.3 kPa) with the proteasome inhibitor MG-132 partially rescue PIP4K2B levels in the cells, indicating a post-transcriptional regulation of this lipid kinase in cells growing in soft conditions (Figure 1C). In order to unveil the mechanosensing role of PIP4K2B we directly silenced this lipid kinase, mimicking its response to soft substrates, and analyzed consequent changes in cell mechanical responses (Figure 1D). PIP4K2B was depleted using pLKO_1 lentiviral vector encoding specific sh_RNAi (sh_2B#1 and sh_2B#2) for this isoform, which had no effects on the expression levels of the other two members of PIP4K family, PIP4K2A and 2C. Empty pLKO_1 vector was used as control (sh_Ctrl) (Figure 1E). Since PIP4K2B seemed to localize in hTERT_RPE1 cells nuclei (Supplementary Figure 1D), we firstly focused on nuclear mechanical properties.

**Figure 1.**
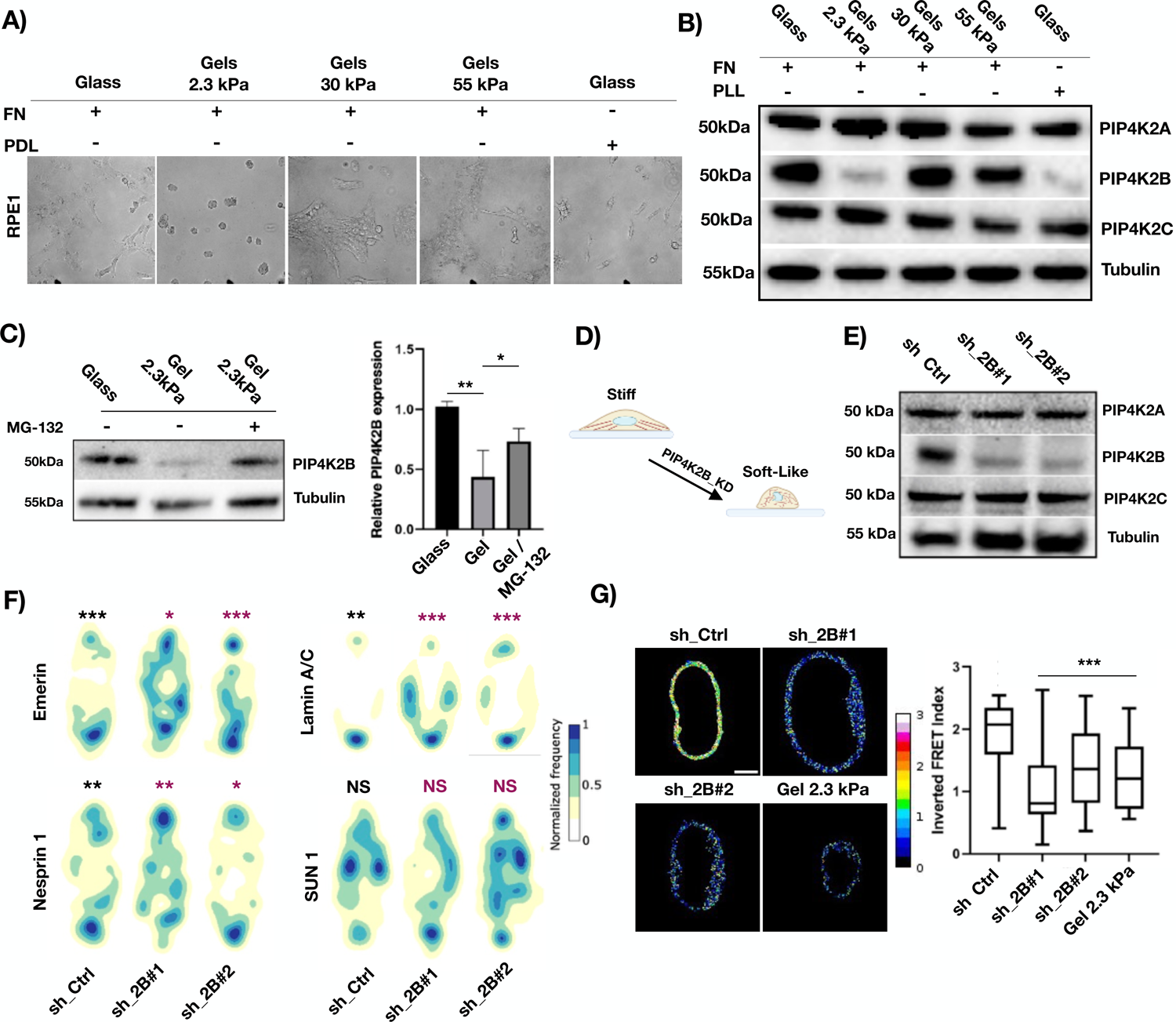
PIP4K2B depletion impairs nuclear mechanical properties. A) hTERT_RPE1 cells were cultured for 24 hours on Fibronectin (FN)- or Poly D-lysine (PDL)-coated glass coverslips, and on FN-coated hydrogels of different stiffness (2.3 kPa / 30 kPa / 55 kPa) (scale bar = 20μm). B) Cell lysates from A) were immunoblotted to analyse PIP4K2A, PIP4K2B and PIP4K2C protein levels. β-Tubulin was used as loading control. C) hTERT_RPE1 cells were cultured as in A). MG-132 (5uM) was added at cell culture medium for 16 hours. Cells growing on Glass, Gel 2.3 kPa and Gel 2.3 kPa+MG-132 were lysed and cell lysates immunoblotted to analyse PIP4K2B protein levels. β-Tubulin was used as loading control. Western Blots were quantified as ratio between intensity of PIP4K2B lanes versus β-Tubulin lanes. Results are reported as average of 5 independent experiments and represented as bar charts +/- standard deviation. D) Graphical representation of biological question about the role of PIP4K2B in driving a soft-like status in the cells. E) hTERT_RPE1 cells were transduced with 2 different sh_RNAi targeting PIP4K2B (sh_2B#1 and sh_2B#2). Western Blotting analysis was performed to analyse the levels of PIP4K2B together with PIP4K2A and PIP4K2C to exclude possible off-target effects of the silencing procedure on other isoforms. Empty pLKO_1 vector was used as Ctrl (sh_Ctrl) F) Nuclear polarity analysis of cells seeded on PLL-g-Peg linear (10μm) patterns: distribution maps of the signal intensity of nuclear proteins analysed through immunofluorescence (Emerin, n=49/51/46, Lamin A/C, n=50/67/50, Nesprin 1, n=41/42/54 and SUN1, n=34/50/49) are reported. At least 34 cells were used for each condition. Black * indicates statistical analysis of the front vs rear accumulation, while purple * indicates statistical analysis of Ctrl cells vs PIP4K2B knock-down cells. G) Nuclear envelope (NE) tension analysis exploiting Mini Nesprin 1 cpst-FRET sensor. Representative nuclei of Ctrl cells (sh_Ctrl, n=47) and cells depleted for PIP4K2B (sh_2B#1/#2, n=51/49) are shown. As internal control, cells seeded on 2.3 kPa gels were used (n=30) (scale bar = 20μm). Data quantification is shown as boxplot chart representing inverted FRET index values (donor/acceptor, the higher the value, the higher the tension) and plotted using Tukey’s method in prism7 software. Statistical analyses were performed on at least 3 independent experiments using unpaired Student’s t test with Welch’s correction or Kolmogorov–Smirnov for maps distribution, *P < 0.05, **P < 0.01, **P < 0.001.

**Figure 2.**
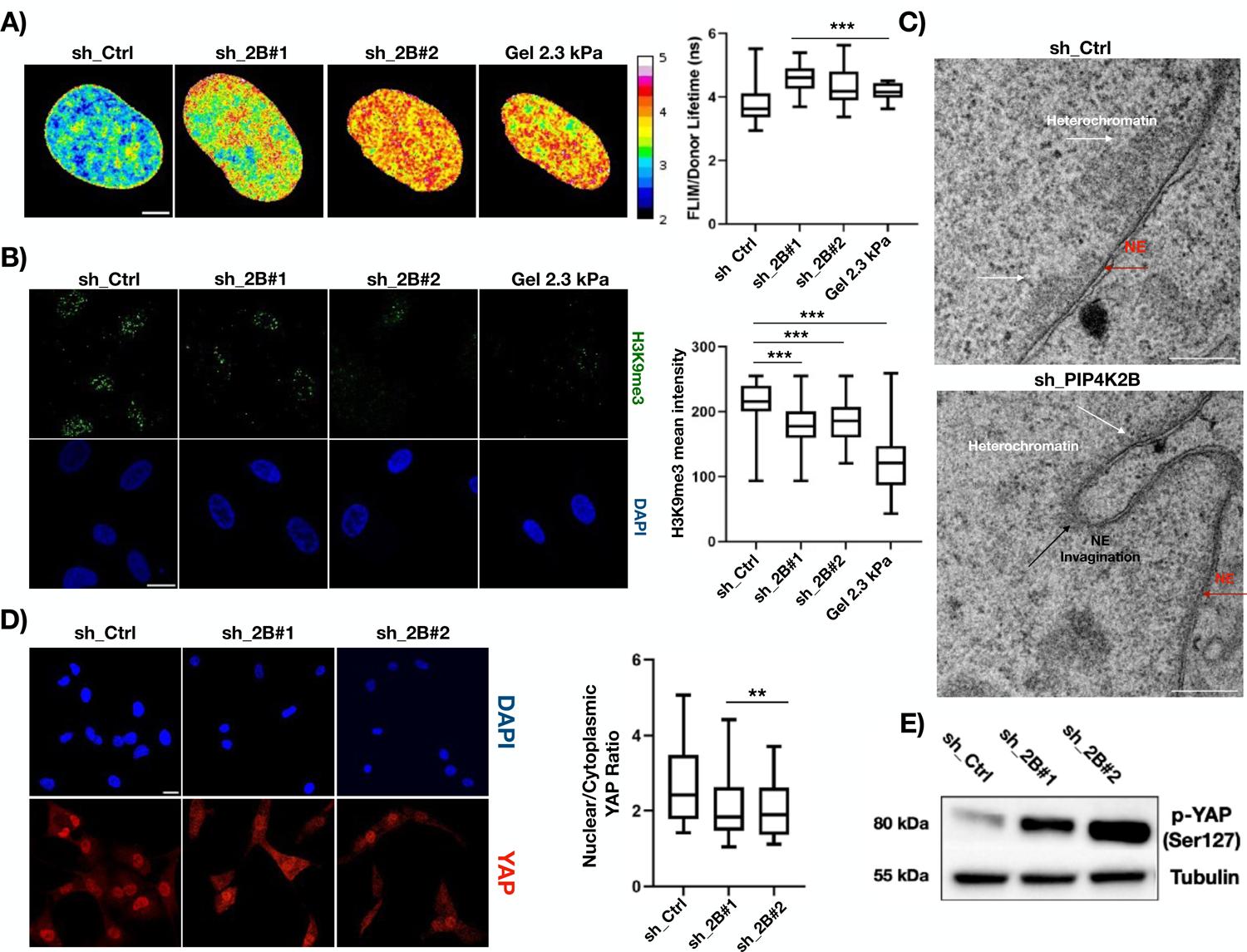
PIP4K2B impacts on chromatin organization altering heterochromatin levels, and drives YAP cytoplasmic localization. A) FLIM-FRET chromatin compaction assay performed transfecting cells with H2B-GFP and H2B-mCherry tagged plasmids. Sample images of Fluorescent lifetime (FLIM-FRET index) of cells transduced with pLKO_1 Ctrl (sh_Ctrl, n=39) and pLKO_1 sh_PIP4K2B (sh_2B#1/#2, n=29/24). As internal control, cells seeded on 2.3 kPa gels were used (n=12). Data are represented as boxplots and overall Lifetime is shown (higher value, lower compaction) (scale bar = 20μm). B) H3K9me3 staining of nuclei of cells transduced with pLKO_1 Ctrl (sh_Ctrl) and pLKO_1 sh_PIP4K2B (sh_2B#1/#2). As internal control, cells seeded on 2.3 kPa gels were used. Data quantification is shown as boxplot representing the ratio between the intensity of single fluorescent dots and cell nuclear area (scale bar = 20μm). C) Transmission electron microscopy (TEM) derived images from routine 60 nm EM section. Colored arrowheads indicate heterochromatin (white), nuclear envelope (NE, black), and NE invaginations (green), respectively (scale bar 1000 ηm). D) Immunofluorescent staining of YAP cellular distribution in sh_Ctrl and sh_2B#1/#2 cells. Quantification of nuclear to cytoplasmic YAP signal ratio are reported as boxplots plotted using Tukey’s method in prism7 software (scale bar = 20μm). E) Western Blotting analysis was performed to analyse the levels of phosphorylated-YAP (Ser-127) in sh_Ctrl and sh_2B#1/#2 cells. Statistical analyses were performed using unpaired Student’s t test with Welch’s correction, *P < 0.05, **P < 0.01, **P < 0.001. At least 50 cells were analysed for IF experiment if not indicated, and data generated come from at least three independent replicates.

**Figure 3.**
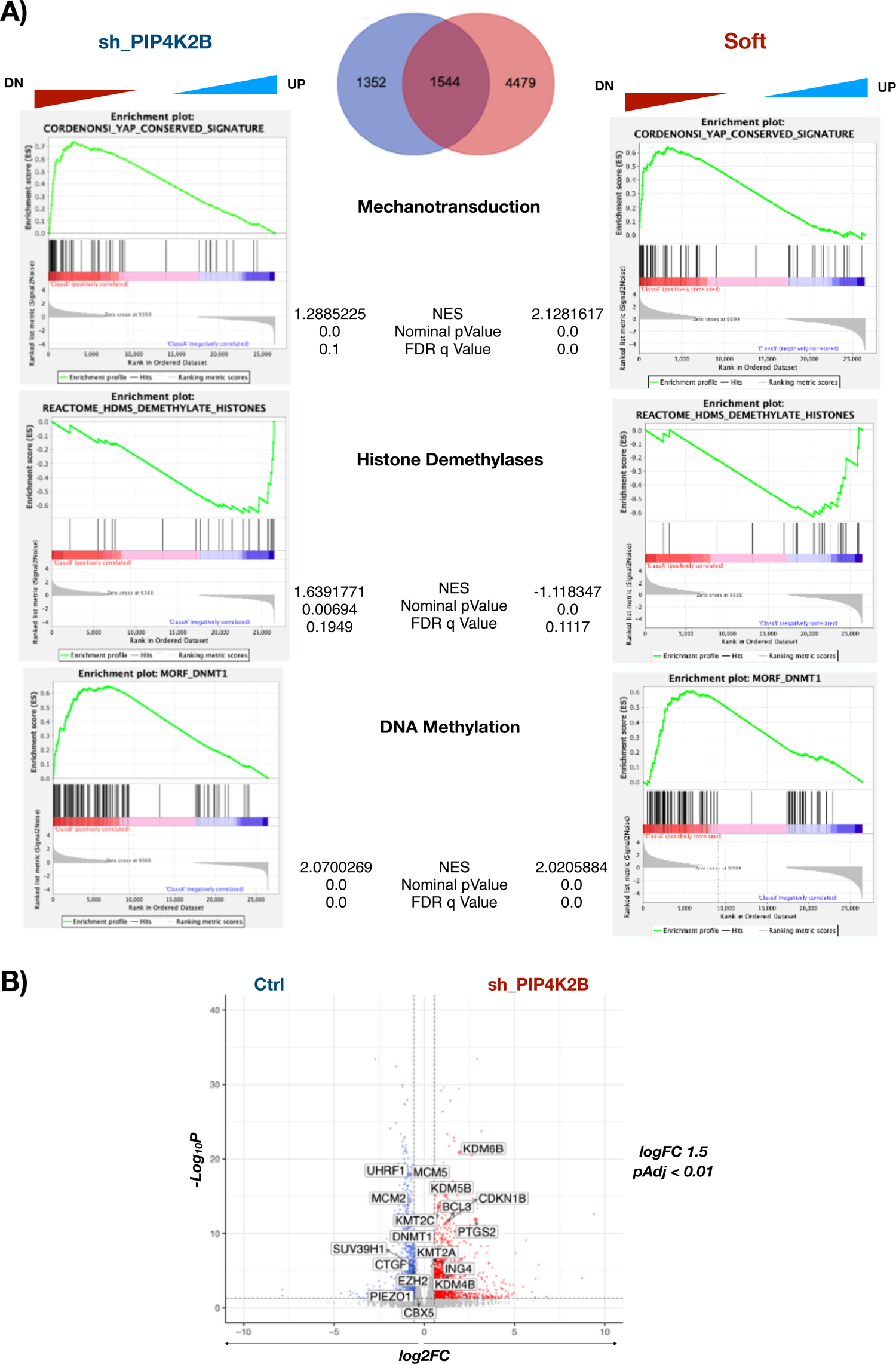
PIP4K2B depletion rewires hTERT_RPE1 cells transcriptomic profile through alteration in mechanotransduction and chromatin remodelling genes. A) Gene expression data generated through RNA-seq comparing Ctrl vs sh_PIP4K2B and Ctrl vs Soft were used for GSEA analyses to extract biological knowledge. False Discovery Rate (FDR) Normalised Enrichment Score (NES) are reported. Venn-Diagram show number of DEGenes in sh_PIP4K2B and Soft conditions if compared to Ctrl. B) Volcano plot representing DEGenes in sh_PIP4K2B vs Ctrl, with highlight of genes involved in chromatin modifications.

### PIP4K2B depletion impacts on nuclear polarization

One of the most important features of animal cells is their capacity to acquire an asymmetric geometry (22, 23). This phenomenon is termed cell polarity, and is deeply connected to substrate stiffness (22, 23). In fact, epithelial cells growing on stiff surfaces are characterized by a well-defined front-rear polarization of their internal organelles, which is lost in cells seeded on soft-like substrates. Cell polarization can be transduced from the cytoplasm onto the nucleus through the connection between nuclear envelope (NE) and cytoskeleton, mediated by the Linker of Nucleoskeleton and Cytoskeleton (LINC) complex (24). We and others showed that direct alteration of proteins of the LINC complex leads to impaired force transduction and defects in front-rear cell polarity transmission to the nucleus (nuclear polarity) (24, 25). In order to investigate if nuclear polarity was altered in cells depleted for PIP4K2B, we plated hTERT_RPE1 cells on FN-coated micro-patterned lines of 10 μm in width, as previously described (24). This allowed the appearance of a clear front-rear polarity in the cells, assesed by Golgi staining used as marker of cell front (Supplementary Figure 2A). We then silenced PIP4K2B and analyzed by immunofluorescence the distribution of NE components. Remarkably, we found that Lamin A/C, Emerin and Nesprin 1 were strongly delocalized upon PIP4K2B depletion (Figure 1F and Supplementary Figure 2B). This variation was not due to changes in the expression of these protein (Supplementary Figure 2C). In addition, we investigated how PtdIns5P, the lipid substrate of PIP4K2B signalling, could localize in polarised cells (Supplementary Figure 2D and 2E). Since no antibody is available to stain for this very rare phosphoinositide, we took advantage of a well-recognized probe for its detection built using the PHD finger of the nuclear protein ING2 and tagged with GFP (26–28). We then silenced PIP4K2B and transfected the cells to allow ING2-PHD_GFP expression. Remarkably, PtdIns5P preferentially localizes at the front of the cell nucleus in control cells seeded onto linear patterns. This localization was completely lost in cells depleted for PIP4K2B. On the opposite, no effects in the distribution of ING2-PHD_GFP were found in the cytoplasm, indicating that knock-down of PIP4K2B mainly impacted on nuclear PtdIns5P localization. Altogether, our findings show that lack of PIP4K2B altered nuclear polarity.

### A novel FRET sensor based on Nesprin 1 shows altered NE tension in cells depleted for PIP4K2B

Considering that lack of PIP4K2B altered the localization of NE components and, possibly, the connection between nucleus and cytoplasm, we investigated the tensional state on the NE(29). For this purpose, we developed an ad hoc fluorescence energy transfer (FRET) sensor using the LINC complex protein Nesprin-1 as backbone, named Mini Nesprin 1, which exploited orientation-based FRET probes (cpst) (30). Nesprins possess a Klarsicht, ANC-1, Syne Homology (KASH) domain at C-Terminus which allows their anchorage to NE through the link with SUN-domain proteins (SUN-proteins binding at perinuclear membrane), and two N-Terminus Calponin domains (CH-CH) for Actin binding (31). Nesprins are then always subjected to a certain degree of force, from relaxed to tensed, depending on the cell mechanical state (32). Our Mini Nesprin-1 FRET sensor can sense the mechanical tension exerted on the NE, i.e. more FRET signal indicates less tension, and viceversa (33, 34). Mini Nesprin 1 showed intense localization at NE and front polarization as wild type protein once expressed in hTERT_RPE1 cells (Supplementary Figure 3A and 3B). Moreover, its expression did not alter cell viability or proliferation (Supplementary Figure 3C, 3D and 3E). We then employed this new tool to decipher if and how depletion of PIP4K2B could impact on NE tension. Cells lacking PIP4K2B were characterized by a strong drop in NE tension (Figure 1G, data presented as Inverted FRET index), in a very similar fashion if compared to cells seeded on a soft substrate (Gel 2.3 kPa). As control of our analysis, we developed a mutant of Mini Nesprin-1, lacking the CH-CH domain. This construct is expected to not react to stretch due to the impaired actin binding and indeed did not show significant changes in FRET signals in cells seeded on either FN- or PDL-coated coverslips (Supplementary Figure 3F, 3G and 3H). These evidences underlie that silencing of PIP4K2B induces a decrease of tension at the NE.

### PIP4K2B depletion impacts on chromatin compaction through alterations of heterochromatin

Alterations in nuclear mechanics are usually followed by changes in chromatin organization (4, 35). In order to study the effects of lack of PIP4K2B on chromatin, we performed chromatin compaction assay exploiting fluorescence energy transfer (FRET)-based fluorescent lifetime imaging microscopy (FLIM) (36). More specifically, cells were transduced to knock-down PIP4K2B and subsequently co-transfected with histone H2B tagged with either -GFP (donor) or -mCherry (acceptor). FRET signals between the two tagged-H2B on separate nucleosomes can be recorded, and changes in donor decay time given by major/minor distance with the acceptor indicate the status of compaction of chromatin, i.e. longer decay time for less compacted chromatin and viceversa. Our findings clearly showed that upon PIP4K2B depletion, the fluorescent lifetime of H2B-GFP resulted increased, indicating a decrease in chromatin compaction (Figure 2A). Strikingly, this chromatin status was similar to the one encountered analyzing cells seeded on soft gels (2.3 kPa), reinforcing the idea that the loss of PIP4K2B is mimicking the cell response to soft substrate. In order to corroborate our findings, we analysed levels of Histone 3 lysine 9 trimethylation (H3K9me3), a well-established marker of heterochromatin, which resulted to be decreased in cells depleted for PIP4K2B, as well as in cells seeded on a soft 2.3kPa gel (Figure 2B) (37). Next, Transmission Electron Microscopy (TEM) analysis also confirmed that this impairment was due to unbalanced levels of heterochromatin (high compaction) and euchromatin (low compaction) in the cell nuclei. We noticed indeed that heterochromatin was almost completely absent in PIP4K2B knock-down cells, in line with a less compacted status of chromatin (Figure 2C and Supplementary Figure 4A, whole images). Interestingly, cells depleted for PIP4K2B also presented strong nuclear invaginations in line with low NE tensional state (Figure 2C and Supplementary Figure 4A, whole images). These evidences suggest that mechano-induced degradation of PIP4K2B drives the rearrangement of cell chromatin state.

### PIP4K2B positively regulates YAP nuclear/cytoplasmic ratio

As PIP4K2B depletion impacts on nuclear mechanics, driving cells in a soft-like status in terms of chromatin compaction and nuclear envelope tensional state, we investigated how this could affect the distribution of the master regulator of mechanotransduction processes YAP (21). We found that strong nuclear localisation of YAP was present in control cells seeded on FN-coated glasses, while nuclear/cytoplasmic ratio was deeply impaired in cells depleted for PIP4K2B (Figure 2D). Consistently, immunoblot analyses showed accumulation of YAP phosphorylation at Ser-127, a well-recognized marker of cytoplasmic retention of YAP (Figure 2E) (38). This is in agreement with the well-established notion that YAP localization is predominantly nuclear in cells growing on stiff substrates, while is more uniform in cells on soft surfaces (21). It has been recently proposed that YAP nuclear import is proportional to forces applied on the nucleus able to dilate nuclear pores sizes (39). Since we observed lower NE tension upon PIP4K2B depletion, we investigated if nuclear transport capacity was altered in PIP4K2B knock-down cells (39). In order to test that, we introduced a GFP tagged with a single nuclear localization signal (NLS) in hTERT_RPE1 cells in which PIP4K2B was silenced and in Ctrl cells. In line with impaired nuclear mechanics in cells lacking PIP4K2B, GFP nuclear staining was strongly impaired in KD cells, suggesting a less active nuclear transport possibly due to the decreased tension at the NE of these nuclei (Supplementary Figure 4B). These findings are in line with recently published data (39), and confirm that proper nuclear mechanics is important for active nuclear import of YAP.

### RNA sequencing unravels PIP4K2B role in mechanotransduction and chromatin remodelling

YAP functions in the nucleus as the master co-transcription factor controlling expression of several genes involved in cell response to mechanical stimuli. Since depletion of PIP4K2B impairs YAP nuclear localisation, we performed RNA-sequencing analysis to investigate how this could affect the transcriptomic landscape of the cells. hTERT_RPE1 cells seeded on FN-coated glass coverslips were then transduced to knock-down PIP4K2B with 3 different sh_RNAi (sh_#1/#2/#3, 1^st^ condition as sh_PIP4K2B), and with empty pLKO_1 vector as controls (2^nd^ condition as Control). In addition, cells transduced with empty pLKO_1 vector and seeded on 2.3 kPa gels were used as soft-seeded cells (3^rd^ condition as Soft). PIP4K2B depletion changed the expression of 2896 genes (pAdj<0.05 and FC >1.5), while cells seeded on soft gels showed 6023 genes changed (pAdj<0.05 and FC >1.5) when compared to control cells (Figure 3A). Interestingly, in line with data showed above, GSEA analysis indicated genes from a YAP conserved signature are more highly enriched in control cells compared to either depletion of sh_PIP4K2B or seeding cells on a soft substrate (Cordenonsi_YAP_Conserved_Signature). This strengthened the concept that lack of PIP4K2B could mimic cell response to soft substrate also at the transcriptomic level (Figure 3A and Supplementary Figure 5A). In particular, once we analysed DEGenes of sh_PIP4K2B vs Control and Soft vs Control, we found an important degree of overlap of genes regulated by PIP4K2B depletion and soft-seeding (Figure 3A). Many DEGenes shared by Soft and sh_PIP4K2B were involved in processes like chromatin remodeling and histone methylation/demethylation (Figure 3A). Moreover, once GO terms were investigated, we found a strongly negative correlation of genes involved in heterochromatin formation in sh_PIP4K2B (Supplementary Figure 5B). This suggests that sh_PIP4K2B cells were characterized by decreased heterochromatin levels, in line with impairment of chromatin compaction and deregulated nuclear mechanics. Since changes in proteins involved in heterochromatin formation and maintenance upon cell mechanical perturbation have been recently indicated as part of the mechanisms controlling heterochromatin-driven nuclear softening (40), we investigated our RNA-seq data, searching for DEGenes involved in heterochromatin remodeling shared by Soft and sh_PIP4K2B conditions. As presented in the Volcano plots and heatmap shown in Figure 3B and Supplementary Figure 5C and 5D, shared DEGenes sets could be divided in 3 different clusters: 1) genes down in Ctrl vs sh_PIP4K2B/Soft (blue); 2) genes up in Ctrl vs sh_PIP4K2B/Soft (red); 3) genes differentially regulated in Soft compared to sh_PIP4K2B (green). We then focused our attention on two genes whose expression was decreased in both sh_PIP4K2B and Soft (cluster 3): SUV39H1 and UHRF1 (Figure 3B and Supplementary Figure 5C and 5D). The first encodes a histone methyltransferase which methylates H3K9me2 into H3K9me3 allowing heterochromatin formation (41). The second encodes for a multidomain nuclear E3-ubiquitin ligase involved in heterochromatin organization, taking part in H3K9 binding of SUV39H1 itself, and in the recruitment of DNA methyltransferases to control DNA methylation (42). Interestingly, we have been able to validate the drop of UHRF1 protein in cells depleted for PIP4K2B while protein levels of SUV39H1 were not altered (Supplementary Figure 6A). Remarkably, UHRF1 protein downregulation was found also in cells seeded on soft substrates, perfectly matching the behavior of PIP4K2B in the same conditions (Supplementary Figure 6B). We recently proposed UHRF1 as a possible target of PIP4K2B signalling, while others showed that its function is tightly connected to its binding to PtdIns5P (43, 44). In addition, UHRF1 gene has been identified as a target of YAP/TEAD transcription factors (45). Then, we decided to investigate how PIP4K2B could be involved in UHRF1 control, and if this was mediated by YAP function.

### PIP4K2B depletion impacts on the chromatin remodeler UHRF1 at post-transcriptional level

Our data showed that silencing of PIP4K2B drove to nuclear envelope tension drop, heterochromatin remodeling, transcriptomic rewiring, possibly due to YAP cytoplasmic retention, and UHRF1 depletion. Then, in order to understand the role of YAP and UHRF1 in the appearance of these phenotypes, we investigated the temporal hierarchy of the events described above. We transduced hTERT_RPE1 cells to silence PIP4K2B and monitored levels of UHRF1 and p-Ser127 YAP for 4 days. Interestingly, we found that upon knock-down of PIP4K2B, UHRF1 levels dropped prior to changes in YAP phosphorylation (Figure 4A). This was confirmed by immunofluorescence staining which showed YAP cytoplasmic retention only at 72h and not at 48h (Supplementary Figure 6C). These data suggest that PIP4K2B controls UHRF1 independently by YAP nuclear localisation. PIP4K depletion usually increases levels of its substrate, PtdIns5P, controlling different nuclear outputs including UHRF1 capacity to bind H3K9me3 (44, 46). In order to mimic the depletion of PIP4K2B, hTERT_RPE1 cells were incubated with exogenous PtdIns5P. More specifically, we starved cells over-night and, next, cultured them in complete medium supplemented with PtdIns5P for 4 hours (+ PI5P). Cells growing without addition of PtdIns5P were used as Control (Ctrl). Strikingly, we found that UHRF1 levels started to decrease just after 4h of PtdIns5P treatment, while YAP was still located mainly in the nuclei of the cells (Figure 4B and 4C). In addition, H3K9me3 levels decreased upon PtdIns5P treatment (Supplementary Figure 6D). This confirms the idea that PIP4K2B impacts primarily on UHRF1 and, subsequently, on YAP signalling. Considering that exogenous PtdIns5P triggered UHRF1 downregulation within 4 hours, we investigated if this could be due to a post-transcriptional regulation of the protein. We then over-expressed UHRF1 in cells previously transduced with sh_Ctrl or sh_PIP4K2B, and we found that also exogenous UHRF1 levels were decreased in cells lacking PIP4K2B. Strikingly, MG-132 treatment could rescue UHRF1 protein in the cells (Supplementary Figure 6E and 6F). Moreover, this was mirrored also in cells seeded in soft-like (PDL) substrates (Supplementary Figure 6G). These data indicated a strong degradation rate of UHRF1 in cells depleted for PIP4K2B, which was confirmed by Ubiquitination assay (Figure 4D).

**Figure 4.**
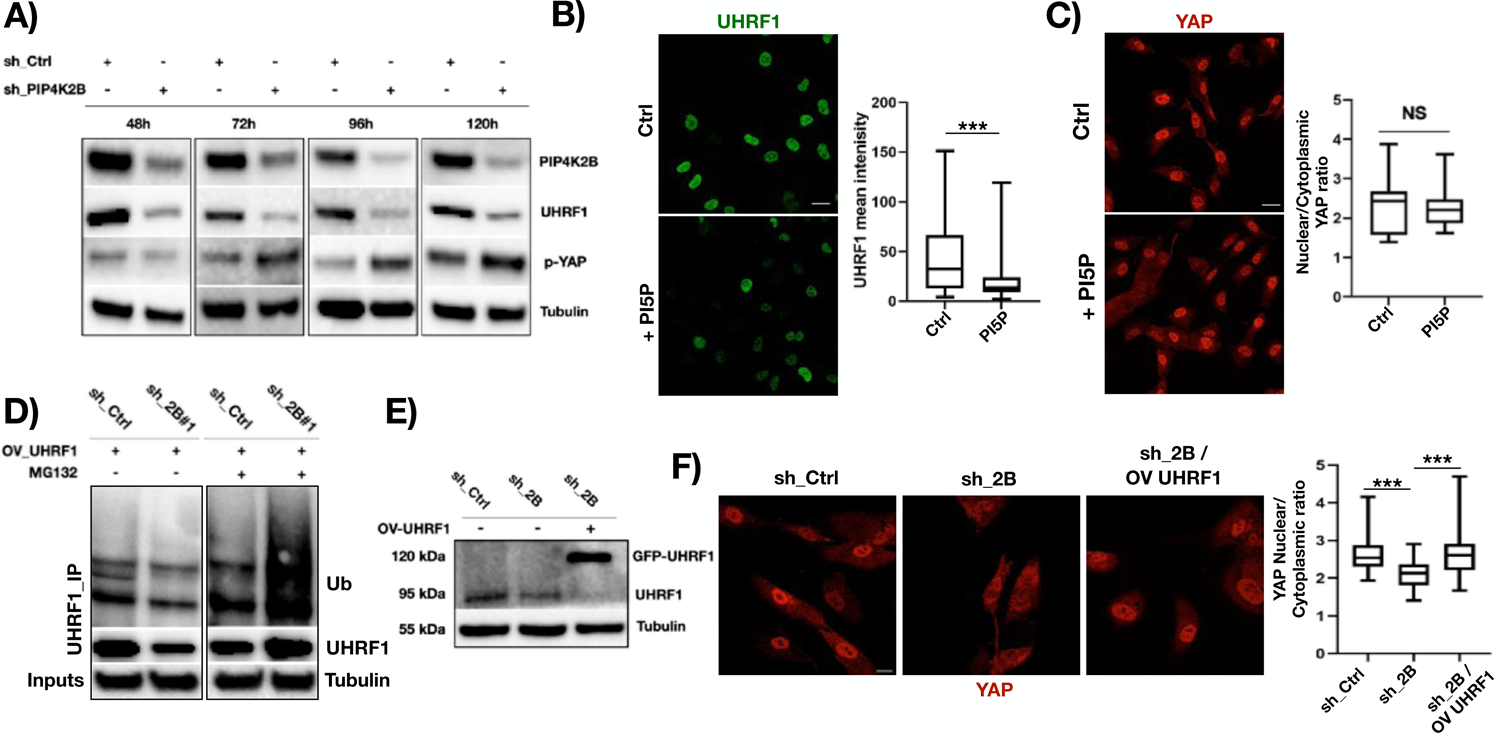
PIP4K2B/UHRF1 signalling controls YAP intracellular localization. A) Time course of the expression of the levels of UHRF1 and phosphorylated-YAP (Ser-127) was assessed by western blotting. WB was performed 48h/72h/96h/120h after that hTERT_RPE1 cells were transduced with sh_Ctrl or sh_2B#1/#2. B) Immunofluorescent staining of UHRF1. Cells were starved for 16 hours, then seeded in complete medium supplemented with PtdIns5P (+PI5P). As control, cells were starved and grown in complete medium (Ctrl) (scale bar = 20μm). Quantification of the mean intensity of UHRF1 is reported. C) Cells were treated as in B) and immunofluorescent staining of YAP cellular distribution was performed (scale bar = 20μm). Quantification of nuclear to cytoplasmic YAP signal ratio is reported. D) Ubiquitination assay showing degradation of UHRF1 in cells transduced to silence PIP4K2B (sh_2B#1/#2). Cells overexpressing UHRF1 were treated with MG-132 (5uM) for 16 hours. As control, cells growing in complete medium were used. UHRF1 was then immunoprecipitated, and western blotting analysis was performed to analyse the levels of Ubiquitinated-UHRF1 (Ub IB). 5% of input samples were used to assess amount of UHRF1 immunoprecipitated, while cell lysates were immunoblotted for Tubulin to check loading conditions. E) Western Blotting analysis of UHRF1 protein expression in cells depleted for PIP4K2B (sh_2B) and concomitantly depleted for PIP4K2B and transduced to overexpress GFP-UHRF1. As control, cells transduced with pLKO_1 vector were used (sh_Ctrl). F) Immunofluorescent staining of YAP localization in cells depleted for PIP4K2B (sh_2B), depleted for PIP4K2B and concomitant overexpression UHRF1 (sh_2B / OV_UHRF1), and control cells (sh_Ctrl) (scale bar = 20μm). Quantification of nuclear to cytoplasmic YAP signal ratio is reported. Data are represented as boxplots plotted using Tukey’s method in prism7 software. Statistical analyses were performed using unpaired Student’s t test with Welch’s correction, *P < 0.05, **P < 0.01, **P < 0.001. At least 50 cells were analysed for every IF experiment, and data generated come from at least two independent replicates.

### UHRF1 depletion phenocopies YAP cytoplasmic retention encountered in cells lacking PIP4K2B

Since PIP4K2B controls UHRF1 protein levels in a YAP independent way, we then investigated if UHRF1 depletion phenocopies PIP4K2B depletion. We thus directly silenced UHRF1 with 2 different sh_RNAs (sh_UHRF1#1/#2) and we analyzed YAP localization in those cells (Supplementary Figure 6H). Interestingly, we found that YAP was strongly localized in the cytoplasm of UHRF1 depleted cells as well as in cells depleted for PIP4K2B (Supplementary Figure 6I). Strikingly, despite the high-rate of UHRF1 degradation in PIP4K2B knock-down cells, once we re-expressed it in these cells, we were able to rescue nuclear YAP localization (Figure 4E and 4F and Supplementary Figure 6J). These data clearly indicate that PIP4K2B controls YAP through UHRF1. On the other hand, the strong decrease in RNA levels of UHRF1 transcript found by RNA sequencing analysis could be explained by a regulatory loop through which YAP cytoplasmic retention drives to impaired YAP-related gene expression of UHRF1, an event which results to be the direct consequence of loss of PIP4K2B and/or UHRF1 themselves.

### PIP4K2B/UHRF1 axis impacts on cytoskeleton and plasma membrane organization

YAP signalling modulates cell mechanics, force transmission and adhesion capacity in the cells, impacting on cell shape, actin cytoskeleton organization and migration (20, 47). In accordance, we found that cells depleted for PIP4K2B or seeded on soft surfaces (gel 2.3 kPa) were characterized by smaller cell area, defects in actin-network, decreased number of stress fibers and focal adhesions (Figure 5A, 5B and Supplementary Figure 7A). This cytoskeletal rewiring was mirrored by decreased phosphorylation levels of focal adhesion kinases (p-FAK) and Myosin Light Chain 2 (p-MLC2), as assessed by western blotting analysis, in cells lacking PIP4K2B (Supplementary Figure 7B). Interestingly, the effect seemed independent by pathways often involved in PIP4K signalling, including PI3K (p-Akt), MAPK (p-ERK) and mTORC1 (p-p70S6K), with a strong downregulation only found in phosphorylation levels of the ribosomoal protein S6 (p-S6) (Supplementary Figure 7B). Again, direct knock-down of UHRF1 phenocopied PIP4K2B depletion in terms of impaired number of focal adhesions (Supplementary Figure 7C). Defects in cytoskeletal organisation and in FA assembly led to altered cytoplasmic rheology and cell capacity to exert forces on the substrate (Figure 5C and 5D). PtdIns5P affects actin polymerization (48), while the pool of PIP4K2B located at the plasma membrane could putatively impact on focal adhesion assembly (49). On the other hand, YAP nuclear extrusion can be controlled by integrin-FA signalling and cytoskeleton remodeling (20). We then decided to confirm the hierarchy of the events triggered by depletion of PIP4K2B and UHRF1 in the cells. We then analysed Focal Adhesion Kinase (FAK) phosphorylation, a marker of FA assembly, by western blotting in cells silenced for PIP4K2B, where UHRF1 levels and YAP phosphorylation have been previously assessed during a time course of events (See Figure 4A). Interestingly, we found that phosphorylation of FAK was lost after 96h of PIP4K2B silencing, an event that occurred only after loss of UHRF1 and YAP phosphorylation, respectively at 48h and 72h (Supplementary Figure 7D, complete WB is presented). As further confirm of this, we investigated FA and actin organization by immunofluorescence (IF). In line with our findings, IF analyses showed impairment of FA and actin cytoskeleton organization in cells depleted of PIP4K2B only after 96h (Supplementary Figure 7E and 7F). Altogether, these evidences indicate a hierarchical model through which PIP4K2B firstly impacts on UHRF1 and, in turn, on FA assembly and cytoskeletal remodeling through an impairment of YAP function in the nucleus.

**Figure 5.**
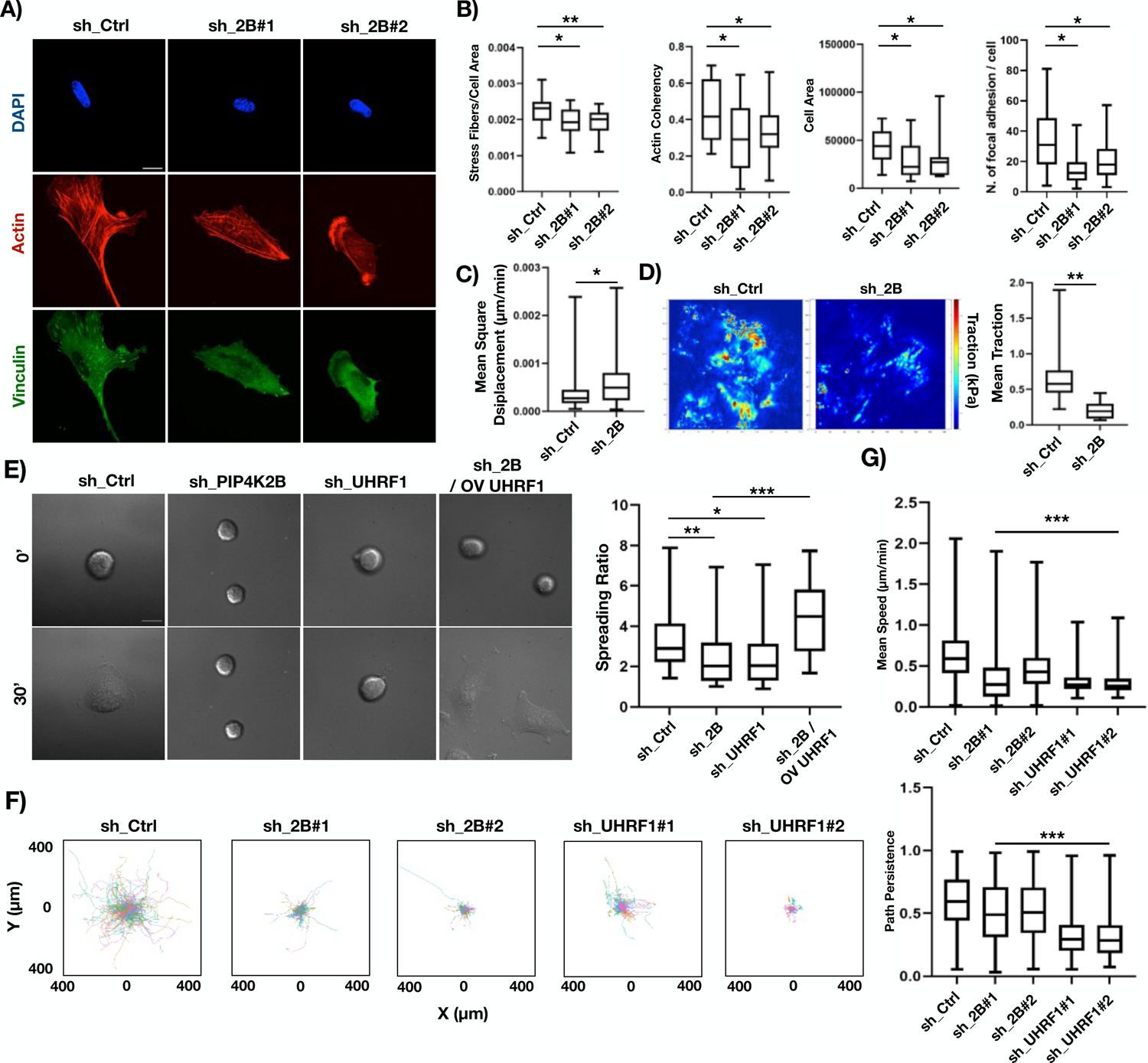
PIP4K2B/UHRF1 signalling controls cytoplasmic organization, impacting on cell capacity to spread and move. A) Immunofluorescent staining of Actin (Phalloidin), Vinculin (Focal Adhesions) and nuclei (DAPI) of cells depleted for PIP4K2B (sh_2B#1/#2). Control cells were transduced with empty pLKO_1 vector (sh_Ctrl) localization. B) Analysis of actin, focal adhesion number, cell area and actin coherency. sh_Ctrl n=18, sh_2B#1 n=25, sh_2B#2 n=25 (scale bar = 10μm). C) Cell Rheology analysis, performed measuring the mean square displacement of microparticles injected in the cell cytoplasm (sh_Ctrl n=40, sh_2B n=40). D) Traction Force Microscopy (TFM) analysis performed on cells seeded for 24 hours on fibronectin-coated silicone (elastic modulus of 5.16 kPa) samples containing QDs. 14 cells were analysed for sh_Ctrl, 25 for sh_PIP4K2B. E) Representative images of cell spreading performed on cells seeded on fibronectin-coated glass coverslips taken at Time 0’ (cell attachment to the substrate) and at Time 30’ (cells in active spreading) (scale bar = 20μm). Movies are available in supplementary material section. Cells were transduced to silence PIP4K2B (sh_PIP4K2B, n=55) or UHRF1 (sh_UHRF1, n=30), and to concomitantly overexpress UHRF1 and silence PIP4K2B (n=27). Control cells were transduced with empty pLKO_1 vector (sh_Ctrl, n=35). F/G) 2D cell motility analysis of cells seeded on fibronectin-coated glass coverslips. Movies are available in supplementary material section. F) Plots representing trajectories of cells depleted for PIP4K2B or UHRF1, and Control cells. G) Data quantification of mean speed and path persistence of the cells. Data are represented as boxplots plotted using Tukey’s method in prism7 software. Statistical analyses were performed using unpaired Student’s t test with Welch’s correction, *P < 0.05, **P < 0.01, **P < 0.001. Data are representative of at least two independent replicates.

### PIP4K2B/UHRF1 signalling controls cells spreading and motility

Since depletion of PIP4K2B or UHRF1 deeply impacted cell mechanics, we investigated how this could be reflected in the motile ability of the cells. We first performed cell spreading analysis. This assay allows evaluation of first phases of cytoskeletal organization, from the initial attachment of the cells to the substrate, followed by quick increase of cell area driven by Arp2/3-dependent actin polymerization, to activation of myosin-dependent contractility (50). Cells transduced to silence PIP4K2B or UHRF1 were allowed to spread on FN-coated glass coverslips. Analyses of the cell area were performed at time 0 (T0’, initial attachment of the cells to the substrate), and after 30 minutes (T30’). The ratio between T30’ vs T0’ cell area clearly showed that cells depleted of PIP4K2B or UHRF1 spread much slower if compared to control cells (Figure 4E). Strikingly, once UHRF1 was re-expressed into cells lacking PIP4K2B, these cells could completely rescue their spreading capability (Figure 5E). Next, we also performed 2D random motility assay, exploiting cells already attached and completely spread on the substrate, to analyze their capacity to move. Cells lacking PIP4K2B or UHRF1 were seeded over-night on FN-coated glass coverslips, and allowed to attach and spread. The day after, time lapse video analyses were performed for at least 18h. In cells depleted for PIP4K2B (with 4 different sh_RNAi) or UHRF1 cell migration capacity was severely impaired (Figure 5F, 5G and Supplementary Figure 8A, 8B and 8C).

### PIP4K inhibition through small molecular compounds mimics effects of PIP4K2B silencing

Due to the increasing interest on the role of PIP4K signalling in cancer, several inhibitors have been developed. We decided to exploit recently synthesized compounds named A-131 and THZ-P1_2 to understand if pharmacological inhibition of these lipid kinases could recapitulate the depletion of PIP4K2B by sh_RNAi (51, 52). We then treated hTERT_RPE1 cells for 24 hours with both these compounds, and we found that, as well as for the knock-down, inhibition of PIP4K led to decreased NE tension and chromatin decompaction (Figure 6A and 6B). This was followed by YAP cytoplasmic retention in treated cells (Figure 5C), and by a strong decrease in cell capacity to move (Figure 5D and Supplementary Figure 9).

**Figure 6.**
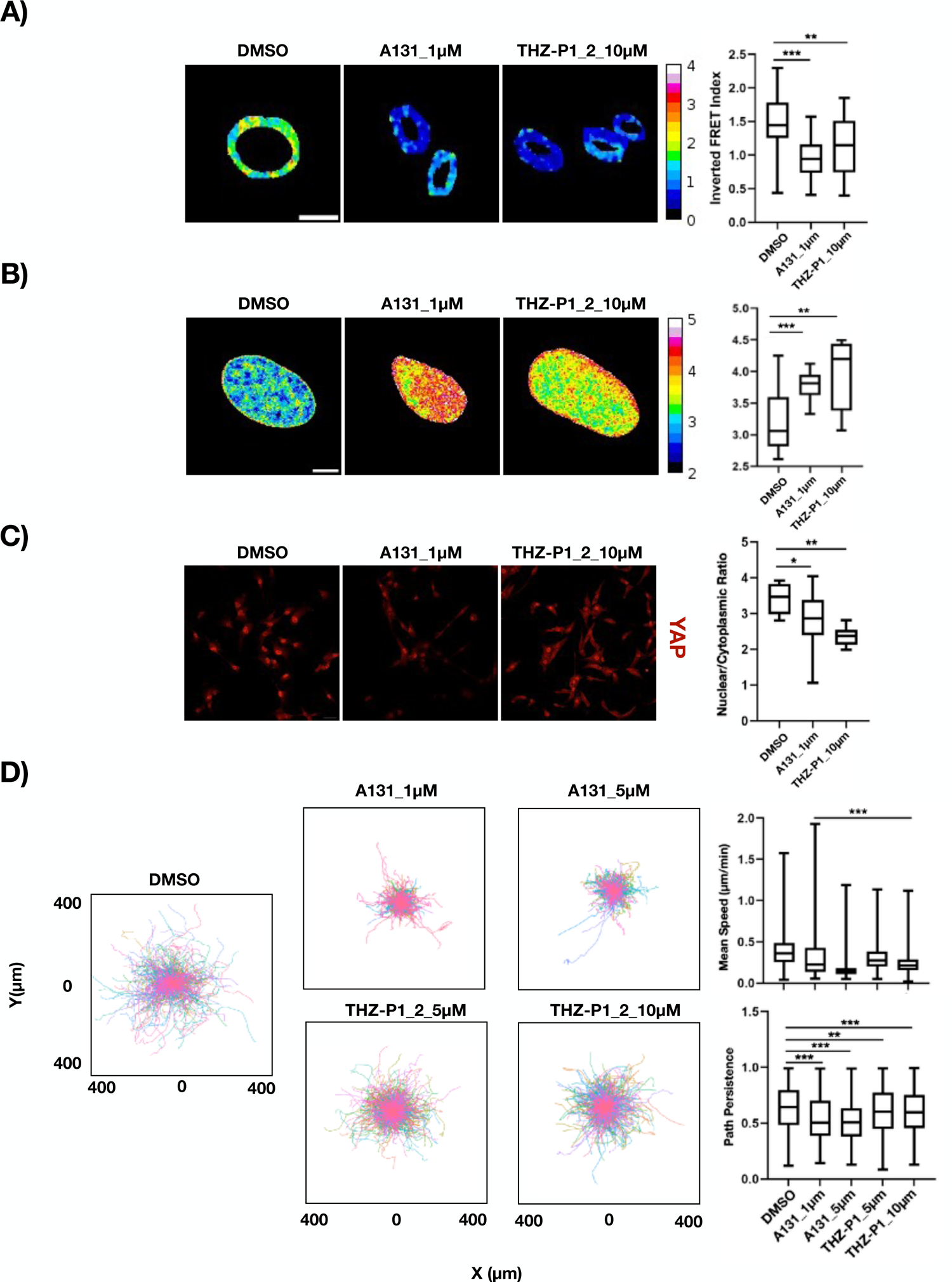
Pharmacological inhibition of PIP4K signalling phenocopies cell softening and defects in motility. A) FRET analysis of NE tensional state exploiting Mini Nesprin 1 cpst FRET sensor. Cells were treated with PIP4K inhibitors A131 (1μM, n=37) and THZ-P1_2 (10μM, n=43) for 24 hours. DMSO was used as vehicle control (DMSO, n=41) (scale bar = 10μm). Data quantification is shown as boxplot chart representing inverted FRET index values (donor/acceptor, the higher the value, the higher the tension). B) FLIM-FRET chromatin compaction assay performed transfecting cells with H2B-GFP and H2B-mCherry tagged plasmids (scale bar = 10μm). Sample images of Fluorescent lifetime (FLIM-FRET index) of cells treated with PIP4K inhibitors as in A). Data are represented as boxplots and overall Lifetime is shown (higher value, lower compaction). PIP4K inhibitor A131 (1μM) n=9, THZ-P1_2 (10μM) n=9, DMSO, n=14. C) Immunofluorescent staining of YAP localization in cells treated as in A). Quantification of nuclear to cytoplasmic YAP signal ratio is reported (scale bar = 20μm). D) 2D cell motility analysis of cells seeded on fibronectin-coated glass coverslips. Movies are available in supplementary material section. Data quantification of trajectories and mean speed/path persistence is shown. Data are represented as boxplots plotted using Tukey’s method in prism7 software. Statistical analyses were performed using unpaired Student’s t test with Welch’s correction, *P < 0.05, **P < 0.01, **P < 0.001. Experiments were repeated three times.

## Discussion

Our study showed that the lipid kinase PIP4K2B functions as a mechanosensor and controls nuclear mechanical properties, impacting on NE and chromatin state. Moreover, we elucidated a mechanism through which PIP4K2B signalling alters levels of the chromatin remodeller UHRF1, inducing nuclear extrusion of YAP and, consequently, rewires cytoskeleton, finally impacting on cell motility.

Cells react to substrate stiffness and timely adapt to the mechanical properties of the microenvironment (53, 54). Here, we showed that cells growing on soft or soft-like substrates (Gel 2.3 kPa and PDL-coated glass) showed a very strong decrease in protein levels of PIP4K2B. Interestingly, the other two members of PIP4K family, PIP4K2A and PIP4K2C, were not altered. In cells growing on soft substrates YAP is mainly localized in the cytoplasm, an event which impairs YAP nuclear signalling impacting on genes involved in cell motility, FA assembly and proliferation (20, 47). However, mRNA levels of PIP4K2B were not altered, indicating that PIP4K2B decrease was not directly connected to YAP function as co-transcription factor. Moreover, inhibiting proteasome-mediated degradation using MG-132 partially rescued PIP4K2B protein levels, suggesting a role for proteasome mediated degradation of PIP4K2B in response to seeding on soft substrates. Inhibition of this lipid kinase can occur through phosphorylation at Ser-326 due to stress related protein kinase p-38 or at Thr-322 by a not identified kinase (55). Previous studies have shown that PIP4K2B can be ubiquitinated by the CUL3 complex (8,) although this did not lead to degradation of the protein. Future works is needed to shed light on mechanisms controlling PIP4K2B expression and stability.

PIP4K2B is highly localized in the nucleus, where it impacts on different nuclear outcomes, including chromatin remodelling and protein-histone binding, through the control of the levels of the two lipid messengers PtdIns5P and PtdIns(4, 5)P2 (7, 9). PIP4K2B resulted strongly accumulated in hTERT_RPE1 cell nuclei, and its depletion strongly affected NE protein localization, altering nuclear polarity. In addition, PIP4K2B depletion changed polarization of the nuclear pool of PtdIns5P, with no effects on the cytoplasmic one, suggesting that PIP4K2B could control its direct substrate mainly at the nuclear compartment. Silencing PIP4K2B also decreased NE tension, which was analyzed with a FRET sensor based on Nesprin 1, named Mini Nesprin 1. In line with previous works, impaired NE tension was also found in cells seeded on soft substrates. Changes in NE tensional state can be due to cytoplasmic forces exerted on the outer nuclear membrane (ONM) through the LINC complex, alterations in the connections between inner nuclear membrane (INM) with chromatin via the lamin meshwork and unbalanced levels of euchromatin and heterochromatin (29). Cells depleted for PIP4K2B showed less compacted chromatin, due to loss of heterochromatin, as indicated by decreased levels of H3K9me3 in the nuclei and by TEM images. These data suggest that the main cause of impaired NE tension found in cells lacking PIP4K2B could be due to a re-arrangement in chromatin organization, and are in line with recent findings showing that alterations in heterochromatin formation through decrease in H3K9me3 levels finally induces nuclear softening (40).

Mechanoresponse to soft-substrate is mainly controlled by retention of YAP in the cytoplasm (20, 56). In line with this, lack of PIP4K2B was followed by YAP nuclear extrusion. Recent findings show how forces acting at the NE, changing nuclear pores size, modulate active nuclear import of small proteins, finally ruling YAP nuclear translocation (39). Our data further support this concept. In fact, cells depleted of PIP4K2B showed altered nuclear import activity, as they were unable to strongly accumulate in their nuclear compartments a GFP tagged with a single NLS.

RNA-seq analysis revealed a high overlap of DEGenes found in PIP4K2B knock-down and in cells seeded on soft surfaces, suggesting that lack of PIP4K2B drives similar transcriptomic rewiring found in cells growing on soft substrates. In fact, once we investigated GSEA datasets related to YAP signalling (GSEA_Cordenonsi YAP Conserved Signature), this was negatively correlated with PIP4K2B silencing. PIP4K2B depleted cells were characterized by impairment in several datasets related to mechanostransduction pathways, in a very similar fashion than cells seeded on soft substrates. These data highlighted the importance of PIP4K2B/PtdIns5P signalling in the control of epigenetic and transcriptional outputs, in response to different substrate stiffness.

In order to understand the mechanosensing role of PIP4K2B, we investigated our RNA-seq data searching DEGenes putatively involved in changes in chromatin organisation. As previously described, alterations in heterochromatin could drive nuclear softening (40). For example, this can be partially due to changes in levels and functions of histone methyltransferases like SUV39H1, the main responsible for H3K9me2 methylation into H3K9me3, which lead to temporary detachment of chromatin from the Lamin A/C meshwork upon mechanical insults (40). Our data indicated several genes involved in heterochromatin formation and maintenance as similarly expressed in PIP4K2B knock-down cells and in soft-seeded cells. We validated our RNA-seq data at protein level, finding UHRF1 as down regulated at both RNA and protein level. UHRF1 is a multi-domain protein which is involved in maintenance of DNA methylation and in chromatin remodelling (42, 57). As part of its functions, UHRF1 acts as a scaffold protein for SUV39H1, facilitating its binding to H3K9me3 (58). It is reported that cells lacking UHRF1 are characterised by defects in methylation status of DNA, and altered heterochromatin conformation (42, 57, 59). UHRF1 contains a Polybasic Region (PBR) able to bind PtdIns5P (44). The binding with this lipid second messenger allosterically regulates UHRF1 protein folding and its histone binding capability. In addition, we recently found that UHRF1 could be a target of PIP4K2B signalling, and others showed UHRF1 gene expression is dependent by YAP/TEAD binding to its promoter (43, 45). Our findings indicate that UHRF1 is targeted by PIP4K2B at post-transcriptional level. PIP4K2B lack drives to UHRF1 degradation, an event which seems to be connected to PtdIns5P, and, consequently to PIP4K2B signalling. How this happens remains to be elucidated. Since previous works suggested that PtdIns5P accumulation could drive to enhanced ubiquitylation of SPOP (speckle-type POZ domain protein) – Cullin3 (Cul3, E3 ubiquitin ligase) substrates (8), it is tempting to speculate that depletion of PIP4K2B may drive UHRF1 degradation through this pathway. On the other hand, our data clearly indicate that this mechanism works independently by YAP function as co-transcription factor. In fact, UHRF1 drop in cells depleted for PIP4K2B occurs prior than YAP phosphorylation and nuclear extrusion. Remarkably, direct silencing of UHRF1 phenocopied PIP4K2B knock-down in terms of YAP cytoplasmic accumulation, and UHRF1 overexpression in cells depleted for PIP4K2B could restore YAP localization in the nuclei. These data suggest a hierarchical series of events which lead to impairment of YAP signalling: first, decreased PIP4K2B levels drive UHRF1 degradation; next, lack of PIP4K2B/UHRF1, inducing chromatin remodelling, impacts on YAP localization; finally, YAP nuclear extrusion is in turn possibly responsible for the subsequent decreased mRNA levels of UHRF1.

Chromatin condensation is necessary for cell motility (60), while YAP signalling controls the expression of genes involved in focal adhesion assembly and cytoskeletal organisation (20). As expected, depletion of either PIP4K2B or UHRF1 drove to strong impairment in focal adhesion number and actin organisation, which was followed by alterations in the capacity of the cells to exert forces on the substrates, spread and properly move. Our data also pointed at a precise hierarchy of the events triggered by depletion of PIP4K2B. Since this lipid kinase is also present in other cell compartments, putatively impacting on cytoskeletal and plasma membrane structural components, which, once altered, can lead to deregulated YAP nuclear/cytoplasmic ratio, we evaluated which was the first target of PIP4K2B silencing (49, 56). Remarkably, due to PIP4K2B lack, UHRF1 down regulation appeared evident at first, followed by YAP nuclear extrusion and, finally, to a rearrangement of focal adhesion assembly and cytoskeletal organisation.

PIP4Ks are highly druggable lipid kinases recently proposed as possible target for the treatment of different cancer types. During the last years, the rising interest in these enzymes led to the synthesis of different inhibitors targeting this family of phosphotransferases. Here, we took advantage of two compounds, named A131 and THZ-P1_2 (51, 52), in order to understand if direct inhibition of the kinase activity of PIP4K could recapitulate the phenotypes appeared upon PIP4K2B silencing. Interestingly, both the compounds altered cell mechanics, finally impacting on cell capacity to migrate, similarly to cells depleted for PIP4K2B.

In conclusion, our studies illustrate the role of PIP4K2B as a mechanosensor. In fact, PIP4K2B depletion alters nuclear mechanics and impact on chromatin organization, through a pathway involving UHRF1, finally altering YAP signalling and cell motility.

Since UHRF1 is highly expressed in cancer, while YAP signalling is known to trigger cancer initiation and growth of solid tumours, our study emphasises the possible benefits given by small compounds interventions targeting PIP4K as treatment for specific YAP-addicted cancer types.

## Materials and methods

### Cell culture

hTERT_RPE1 cells were cultured in DMEM/F12 (Sigma Aldrich) medium supplemented with Penicillin/Streptomycin (1X) and Glutammine (1X). HEK293T, MEF and Hela cells were cultured in DMEM medium (Sigma Aldrich) supplemented with Penicilin/Streptomycin and Glutamine. Inhibition of PIP4K was obtained treating cells with THZ-P1_2 (5/10μM) or PIP4K-in-A131 (1/5μM, MedChem. express) for 24 hours. PI5P (20μM) treatment was performed in cells growing in complete medium for 4h, after a period of 16h of serum starvation. MG-132 treatment (5μM) was performed over-night in growing cells.

### Nuclear Envelope tension analysis and synthesis of a novel cpst-FRET sensor

Nuclear Envelope (NE) tension was analysed using a novel cpst (30) based FRET sensor built using Nesprin 1 protein as backbone, and named Mini-Nesprin 1. N-terminus Nesprin 1 (1521bp, aa 1-507) and C-terminus (1422bp, aa 8325-8797) coding sequences were synthesized exploiting IDT gBlocks Gene Fragments Technology. Briefly, gBlock 1 encoding N-terminus Calponin domains (CH-CH) of giant Nesprin 1 was cloned into Spectrin-cpstFRET (plasmid#61109, Addgene) cut with AgeI and ScaI renstriction enzymes (New England Biosciences). Then, cpst FRET probe from Addgene plasmid #61109 was PCR amplified and cloned between ScaI and KpnI restriction sites. Subsequently, gBlock 2 encoding C-terminus KASH domain was cloned between KpnI and MfeI restriction sites. A truncated version of the FRET sensor (CH-mutant) used as control in the setting-up of the experiments was obtained removing the first 751bp from the original plasmid in order to brake CH domain and, then, impairing Actin binding of Mini_Nesprin 1. Snapgene maps of the constructs are available as Supplementary Materials, and graphical presentation of Mini_Nesprin 1 versions is reported in Supplementary Figure 3F and 3G. Image acquisition was performed using a Leica TCS SP8 confocal microscope with microscope with Argon light laser as excitation source tuned at 458 nm and HC PL APO CS2 ×40/1.40 oil-immersion objective. Emission signals were captured every 10nm starting from 460nm up to 600nm, and inverted FRET index calculated as the ratio between cpCerulean (∼480nm) and cpVenus (∼530nm) peaks. Data were analyzed using in-house ImageJ macro.

### Chromatin compaction assay and FLIM analysis

Chromatin compaction analysis was performed as originally reported (36). Briefly, cells were transfected using Neon electroporation system (Thermo Fischer) with pBabe_H2B-mCherry and pCDNA_H2B_GFP plasmids. Next, transfected cells were seeded on glass coverslips coated with Fibronectin (+/-hydrogels) and analysed using a Leica TCS SP8 confocal microscope with White light laser as excitation source tuned at 488 nm and HC PL APO CS2 ×40/1.40 oil-immersion objective, everything managed by Leica Application Suite X software, ver. 3.5.2.18963. For the lifetime measurements, the above system was implemented with PicoQuant Pico Harp 300 TCSPC module and picosecond event timer, managed by PicoQuant software (SymPho Time 64, ver. 2.4). Data were imported and analyzed using in-house ImageJ macro.

### Traction Force Microscopy

Red fluorescent quantum dots (QDs) were deposited on silicone samples into confocal monocrystalline triangular arrays as described here (61). In the experiment of TFM, hTERT_RPE1 cells were transduced and 4 days later seeded for 24 hours on fibronectin-coated silicone (elastic modulus of 5.16 kPa) samples containing QDs. Cells were then analysed live using oil immersion 60x objective (Leica Germany) with Nikon Eclipse Ti microscope (Nikon Instruments). Reconstruction of the traction field from a single image of the displaced QDs exploited Cellogram software (61). Displacement field to the QDs’ resting position was inferred, and tractions calculated using the known material properties.

### Micro-Rheology analysis

Particle-tracking microrheology experiments were performed as previously described with minor changes. Carboxyl-modified fluorescent polystyrene particles (0.50 µm diameter, Polyscience) were introduced hTERT_RPE1 cells by using a ballistic gun (Bio-Rad). Helium gas at 900 psi was used to force a macro-carrier disk coated with particles to crash into a stopping screen. The force of collision was transferred to the particles, causing their dissociation from the macro-carrier and the bombardment of cells. Once bombarded, cells were extensively washed with PBS and left to recover for 24 h in culture medium. After recovery, cells were seeded (10^4^ cells·cm^-2^) on 35 mm glass bottom Petri dishes (Ibidi). After 24 h from cell seeding, the motion of nanoparticles embedded inside the cells was recorded for a total of 5 s at 100 frames per second using an inverted fluorescence microscope (Olympus IX81; Olympus) equipped with a 100× oil immersion objective (N.A. = 1.40), plus 1.6× magnification of internal microscope lens, and a Hamamatsu ORCA-Flash 2.8 CMOS camera (Hamamatsu). Experiments were performed under physiological conditions, using a microscope stage incubator (Okolab) to keep cells at 37 °C with 5% CO_2_. The total number of analysed particles was at least 80 from more than 10 cells for both cell lines.

Particle-tracking microrheology allows to have indirect information about the local viscoelastic properties of living cells with a high spatiotemporal resolution, collecting and analysing the Brownian motions of particles embedded in the cytoplasm. By using our self-developed Matlab cose, the point tracking trajectories were generated in two distinct steps: firstly, the beads are detected in each frame, and then the points are linked into trajectories. Each position was determined by intensity measurements through its centroid, and it was compared frame by frame to identify the trajectory for each particle, based on the principle that the closest positions in successive frames belong to the same particle (proximity principle). Once the nanoparticle trajectories had been obtained, mean squared displacements (MSDs) were calculated from the following equation

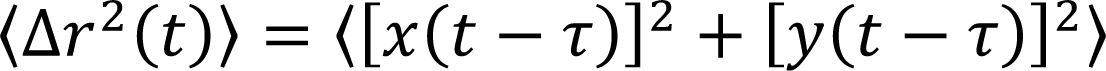

where angular brackets mean time average, τ is the time scale and t is the elapsed time.

### Cell transfection, lentiviral production and cell transduction

Cell transfection was performed using Neon Transfection System (Thermo Fischer). hTERT_RPE1 cells were transfected using the following parameters: Voltage: 1350V, Width: 20ms, Number of pulses: 2×10^6^ cells were transfected with ING2-PHD_GFP (Addgene, #21589) or pCSDEST2_NLS-GFP (Addgene, #67652 doi: 10.1016/j.devcel.2015.10.016.) plasmids. Lentiviral particles were produced as described here (43). Briefly, 3×10^6^ HEK293T cells were transfected with pLKO_1 (encoding sh_RNAi) or pCMV6-AC (Origene #RG217766, encoding GFP-UHRF1) lentiviral plasmids, together with psPax2 and pMDG2 vectors with a ratio of 4μg:2μg:1μg. Polyethylenimine (PEI) was used as transfection reagent. 24h later medium was removed, and virus collection started after 24/48/72h. Viral aliquots were pulled together and filtered with 0.45μm filters. 2×10^5^ cells were transduced with 1ml of fresh virus through 20’ of spinoculation at 2000rpm, then seeded in 6 well plates. sh_RNAi sequences are reported in supplementary informations in Table 4.

### Western Blotting

Cell lysates were quantified analysing absorbance of the samples (595nm) diluted in Bradford (Biorad) at spectrophotometer. They were next separated on bis-Tris sodium dodecyl sulfate– polyacrylamide gel electrophoresis gels, electroblotted to nitrocellulose, blocked in PBS–Tween-20 (0.1%) dried milk (5%), and then incubated with the primary antibodies. After washing, blots were incubated with appropriate secondary antibodies and developed using ECL-Plus (Biorad) at Chemidoc (Biorad). Quantification was performed through Fiji as the ratio between PIP4K2B and Tubulin (housekeeping) signals. A complete list of antibodies exploited in this work is available in table 1 as supplementary material.

**Table 1.** WB antibodies

| Target | Manufacturer | Catalogue Number |
| --- | --- | --- |
| PIP4K2A | Cell Signalling | 5527 |
| PIP4K2B | Cell Signalling | 9694 |
| PIP4K2C | Proteintech | 17077-1-AP |
| Tubulin | Sigma | sc-5724 |
| UHRF1 | Santa Cruz | sc-373750 |
| SUV39H1 | Santa Cruz | sc-23961 |
| p-FAK | Santa Cruz | sc-374668 |
| p-YAP | Cell Signalling | 4911 |
| p-S6_Ser235/236 | Cell Signalling | 2211 |
| p-p70S6K | Cell Signalling | 9205 |
| p-Akt_Ser473 | Cell Signalling | 4060 |
| p-MLC2 | Cell Signalling | 3674 |
| p-ERK1/2 | Cell Signalling | 9101 |
| Lamin A/C | Santa Cruz | sc-7292 |
| Lamin B1 | Abcam | ab16048 |
| Nesprin 1 | Sigma Aldrich | HPA019113 |
| SUN1 | Abcam | ab124770 |
| Emerin | Novocastra | 6044962 |
| Ubiquitin | Santa Cruz | sc-8017 |

### Ubiquitination assay

Cells overexpressing UHRF1 were treated over-night with MG-132 at 5μM as final concentration. Next, cells were lysed with RIPA buffer supplemented with MG-132 1μM, DUB inhibitor PR-619 and beta-Mercaptoethanol. 500μg of cell lysates immunoprecipitated using UHRF1 antibody (Santa Cruz) following manufacturer’s protocol. Protein A/G coated beads were added to the mix, and everything was incubated over-night a 4C in rotation. The next day, beads were washed 5 times with lysis buffer and resuspended with 50ul Loading buffer 4X (Thermo Fischer) plus Reducing Agent 1X. Western blotting was then performed and samples were immunoblot with Ubiquitin antibody. Amount of UHRF1 immunoprecipitated was analysed using 5μl of immunoprecipitated lysates, while inputs were loaded and immunoblotted a part in order to control quality of equal-loading of different samples (using Tubulin as housekeeping).

### RT-qPCR

RNA was extracted with RNA extraction kit (Invitrogen), and quantified at Nanodrop. 100ng of RNA were retrotranscribed into cDNA using a Reverse-Transcription kit (Life Technologies). qRT-PCR was performed using SYBR Green (Life Technologies) to detect expression of PIP4K2B and the GAPDH gene used as housekeeping control (primers sequences reported in Table 3).

### RNA sequencing

For each experimental sample, totRNA was extracted using the RNeasy Prep kit (QIAGEN), and its abundance was measured using Qubit 4.0 and integrity assessed using Agilent Bioanalyzer 2100 using Nano RNA kit (RIN > 8). For each sample, an indexed-fragment library was prepared starting from 500ng totalRNA using Illumina Stranded mRNA Prep ligation kit (Illumina) according to manufacturer’s instructions. Indexed libraries were quantified with Qubit HS DNA kit and controlled for proper size using Agilent Bioanalyzer 2100 High Sensitivity Dna kit prior to normalization and equimolar pooling to perform a multiplexed sequencing run. 5% of Illumina pre-synthesized PhiX library was included in the sequencing mix, to serve as a positive control. Sequencing was performed in Paired End mode (2×75nt) on Illumina NextSeq550Dx instrument, generating on average 60 million PE reads per each library. Reads were aligned to the GRCh38/hg38 assembly human reference genome using the STAR aligner (v 2.6.1d).

Differential gene expression analysis was performed using the Bioconductor package DESeq2 (v 1.30.0) that estimates variance-mean dependence in count data from high-throughput sequencing data and tests for differential expression exploiting a negative binomial distribution-based model. Preranked gene set enrichment analysis (GSEA) for evaluating pathway enrichment in transcriptional data was carried out using the Bioconductor package fgsea (v 1.16.0) and GSEA software taking advantage of the Reactome, KEGG, oncogenic signature and ontology gene sets available from the GSEA Molecular Signatures Database (https://www.gsea-msigdb.org/gsea/msigdb/genesets.jsp?collections).

### Cell motility assay

3×10^5^ hTERT_RPE1 cells were seeded in 6 well plates over-night before live-microscopy analysis. Then, cells were stained with NucBlue Live Cell Reagent (Thermo Fischer) and imaged every 10 minutes for 18 hours using a humidity- and temperature-controlled inverted wide-field microscope within an environmental chamber (ScanR, Olympus). Nuclear tracking and cell motility analysis were performed as described here (24). For this purpose, specific software in C++ with the OpenCV [http://opencv.willowgarage.com/wiki/] and the GSL [http://www.gnu.org/software/gsl/] libraries was developed. The migration analysis was performed by the C++ software coupled with R [www.R-project.org].

### Cell spreading

Cells at different experimental conditions were trypsinized and resuspended in 1x Ringer buffer (150 mM NaCl, 1mM MgCl2, 1mM CaCl2, 20 mM Hepes (pH 7.4), 5 mM KCl and 2g/l glucose).

Suspended cells were seeded on fibronectin-coated glass coverslips (10 μg/ml) directly on the microscope stage at temperature-controlled conditions. Imaging was performed in DIC mode on the Leica AM TIRF MC microscope by a HCX PL APO 63 × /1.47NA oil immersion objective. Cell spreading was followed for 30 minutes up to 1h at a frame rate of 2’. Analysis of spreading capacity was calculated as spreading ration between time 0 and time 30’.

### Transmission Electronic Microscopy (TEM)

Electron microscopic examination was performed as previously,^62^., a detailed description is explained below. EM based Morphological analysis: RPE1 cells grown on MatTek glass-bottom dishes (MatTek Corporation, Cat. P35G-1.5-14-C) for 24 h. Samples were fixed with of 4% paraformaldehyde and 2,5% glutaraldehyde (EMS) mixture in 0.2 M sodium cacodylate pH 7.2 for 2 hat RT, followed by 6 washes in 0.2 sodium cacodylate pH 7.2 at RT. Then cells were incubated in 1:1 mixture of 2% osmium tetraoxide and 3% potassium ferrocyanide for 1 h at RT followed by 6 times rinsing in cacodylate buffer. Then the samples were sequentially treated with 0.3% Thiocarbohydrazide in 0.2 M cacodylate buffer for 10 min and 1% OsO4 in 0.2 M cacodylate buffer (pH 6,9) for 30 min. Then, samples were rinsed with 0.1 M sodium cacodylate (pH 6.9) buffer until all traces of the yellow osmium fixative have been removed, washed in de-ionized water, treated with 1% uranyl acetate in water for 1 h and washed in water again. The samples were subsequently embedded in Epoxy resin at RT and polymerized for at least 72 h in a 60 °C oven. Embedded samples were then sectioned with diamond knife (Diatome) using Leica ultramicrotome. Sections were analyzed with a Tecnai 20 High Voltage EM (FEI) operating at 200 kV.

### Immunofluorescence

Cells were seeded on Fibronectin (20μg/μl) or PDL (0.1mg/ml) coated coverslips (+/- hydrogels) for the time indicated in the experiments. Hydrogel preparation was performed using recipes reported in table 5. Next, they were washed in PBS 1X, fixed with 4% paraformaldehyde for 10’ and washed again 3 times with PBS 1X. Fixed cells were permeabilized with Triton 0.2% for 10’, and incubated for 1h in BSA 0.2%. Primary antibodies staining was performed in a humidity-controlled dark chamber at room temperature for 2h, and removed by 3 times of PBS washing. Secondary antibodies were used in dark at room temperature for 1h, and nuclei stained with 4ʹ,6-diamidino-2-phenylindole (DAPI, Sigma-Aldrich Cat. D8417) for 5 minutes. Samples were prepared using Glycerol as mounting medium and analysed. A complete list of the antibodies employed in this study is available in Table 2 in supplementary materials. Analysis of FA number, Actin organization, and YAP nuclear/cytoplasmic ratio were performed exploiting custom Fiji scripts.

**Table 2.** IF antibodies

| Target | Manufacturer | Catalogue Number |
| --- | --- | --- |
| PIP4K2B | Ref | PIP4K2BP6 |
| Lamin A/C | Santa Cruz | sc-7292 |
| Vinculin | Sigma | V9264 |
| Nesprin 1 | Sigma Aldrich | HPA019113 |
| SUN1 | Abcam | ab124770 |
| Emerin | Novocastra | 6044962 |
| Actin_Phalloidin | Sigma | P2141 |
| H3K9me3 | Cell Signalling | 13969 |
| UHRF1 | Santa Cruz | sc-373750 |

**Table 3.**
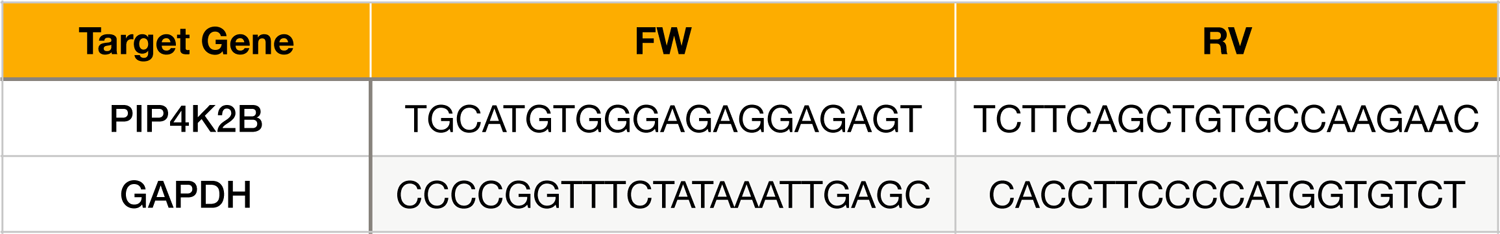
RT-qPCR primers

**Table 4.**
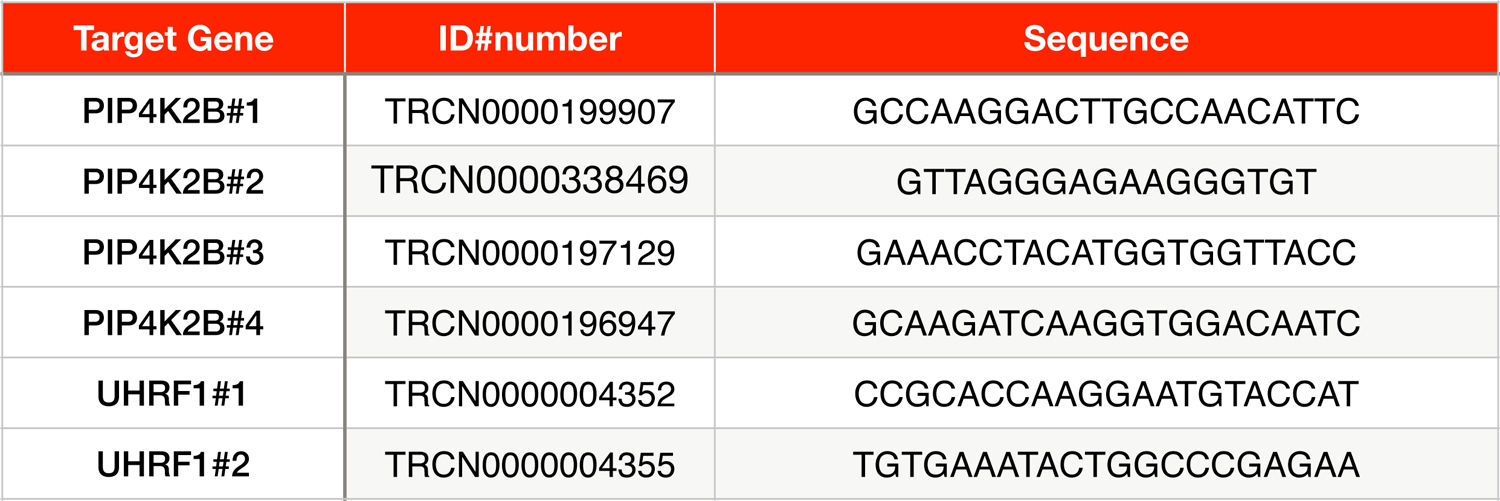
sh_RNAi sequences

**Table 5.** Hydrogel preparation

|  |  |  |  |
| --- | --- | --- | --- |
| **Shear Modulus (Pa)** | **230** | **34263** | **55293** |
| **40% Acrylamide (mL)** | 1.25 | 2.50 | 2.50 |
| **2% Bis-Acrylamide (mL)** | 0.5 | 1.88 | 2.50 |
| **H2O (mL)** | 3.25 | 0.63 | 0.00 |
| **Volume total (mL)** | 5 | 5 | 5 |
| **Stock solution used (Pa)** | **230** | **34263** | **55293** |
| **Stock solution volume (uL)** | 150 | 300 | 300 |
| **TEMED (uL)** | 347 | 197 | 197 |
| **10 % APS (uL)** | 0.7 | 0.7 | 0.7 |
| **H2O (uL)** | 2.5 | 2.5 | 2.5 |
| **Volume Total (uL)** | 500 | 500 | 500 |
| **Final Acrylamide %** | 3 | 12 | 12 |
| **Final Bis-Acrylamide %** | 0.06 | 0.45 | 0.6 |

### Nuclear Polarity analysis

Nuclear polarity analysis was performed as described here (24). Briefly, cells were seeded on fibronectin-coated micro-patterned lines 10 μm. Patterns were fabricated using photolithography. The glass surface of the coverslips was treated and activated with plasma cleaner (Harrick Plasma), then coated with PLL-g-PEG (Surface Solutions GmbH, 0.1 mg/mL in 10 mM HEPES). After washing in PBS 1X, surface was illuminated with deep UV light (UVO Cleaner, Jelight) through a chromium photomask (JD-Photodata). Coverslips were then incubated with Fibronectin (25 μg/ml), and cells seeded over-night. Cell polarization was assessed using Golgi staining marker Giantin (front), and IF was performed for the analysed NE components.

Maps of nuclear polarity were created using a custom Fiji macro () which encounts the top 20% of the signal of the antibodies exploited in the IF. Images were taken every 0.6 μm of focal plane using z-stack function using oil immersion ×40/×60 objective (Leica Germany) at Nikon Eclipse Ti microscope (Nikon Instruments) equipped with the UltraVIEW VoX spinning-disc confocal unit (PerkinElmer) and Velocity software (PerkinElmer).

### Cell Cycle Analysis

Cells growing in complete medium where collected, washed in PBS, and fixed in cold EtOH 70% overnight. Cells were washed with PBS and stained with propidium iodide (Abcam) and analyzed by FACS.

### Cell Apoptosis Analysis

Cells were resuspended in Annexin V resuspension buffer and stained with Annexin V-FITC (BioLegend) for 20 min at room temperature and then directly analyzed by FACS.

### Cell Proliferation

Cells were seeded at the same number in 24 wells (1×10^4^) in complete medium and counted manually for 3 days (24h/48h/72h after seeding).

### Real Time qPCR

RNA was extracted from cells using a commercial kit (Invitrogen). Complementary DNA was synthesized using a commercial kit (Thermo Fischer), and RT-qPCR analysis was performed using SybrGreen (Roche) to detect PIP4K2B expression. GAPDH was used as housekeeping gene.

## Statistical analysis

Statistical analysis war performed using Unpaired t-test with Welch’s Correction, and two-sided Kolmogorov–Smirnov test for nuclear polarity maps distribution. Every experiment was performed at least 2 times, independently. Statistical significative data are considered data with a pValue < 0.05.

## Supporting information

Cell_spreading_Video

Cell_spreading_Video

Cell_spreading_Video

Cell_spreading_Video

2D_Cell_Motility_Video

2D_Cell_Motility_Video

2D_Cell_Motility_Video

2D_Cell_Motility_Video

2D_Cell_Motility_Video

## Acknowledgments

This project was supported by Italian Association for Cancer Research (AIRC) Investigator Grant (Paolo Maiuri, #24976) and individual fellowship (Fabrizio A. Pennacchio, #23966). Alessandro Poli’s work was founded by Fondazione Umberto Veronesi Post-doctoral fellowships (#000359) and Short-EMBO Fellowship (#8386). The authors would like to thank IFOM Imaging Facility for help with performing experiments. We would like to thank Dr. Carlos Nino for help in setting the Ubiquitination assay.

## Supplementary Figures legends

**Supplementary Figure 1.**
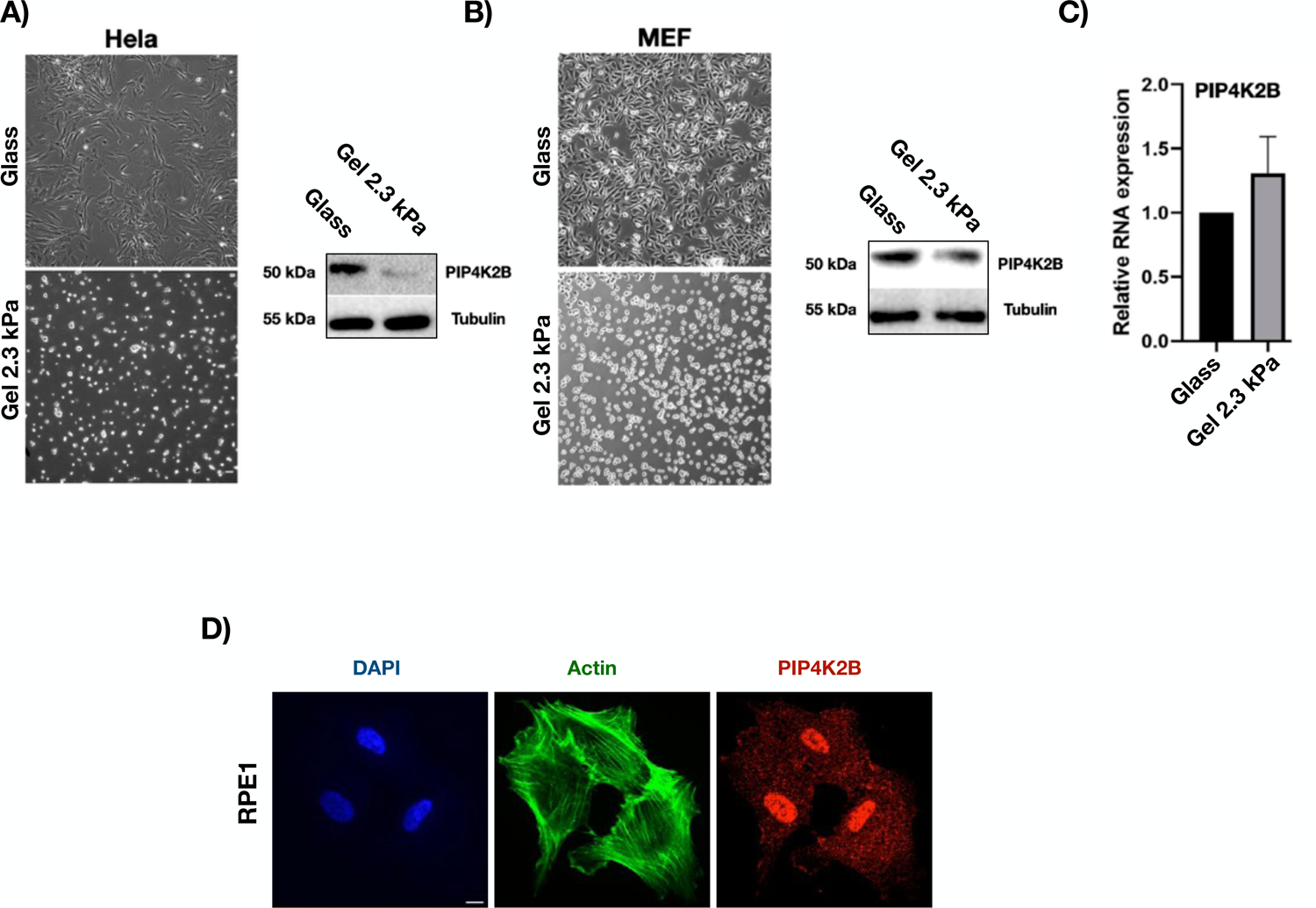
A/B) Hela and MEF cells were grown on FN-coated glass (stiff) or elastic surface (Gel 2.3kPa, soft) for 24h, then lysed (scale bar = 20μm). Protein lysates were then immunoblotted to analyse PIP4K2B expression. β-Tubulin was used as loading control. C) RT-qPCR analysis of mRNA levels in hTERT_RPE1 cells grown as in A). GAPDH was used as housekeeping gene. Data are shown as Log2FoldChange. D) Immunofluorescence staining of hTERT_RPE1 cells for PIP4K2B, Actin (Phalloidin) and nuclei (DAPI) (scale bar = 10μm).

**Supplementary Figure 2.**
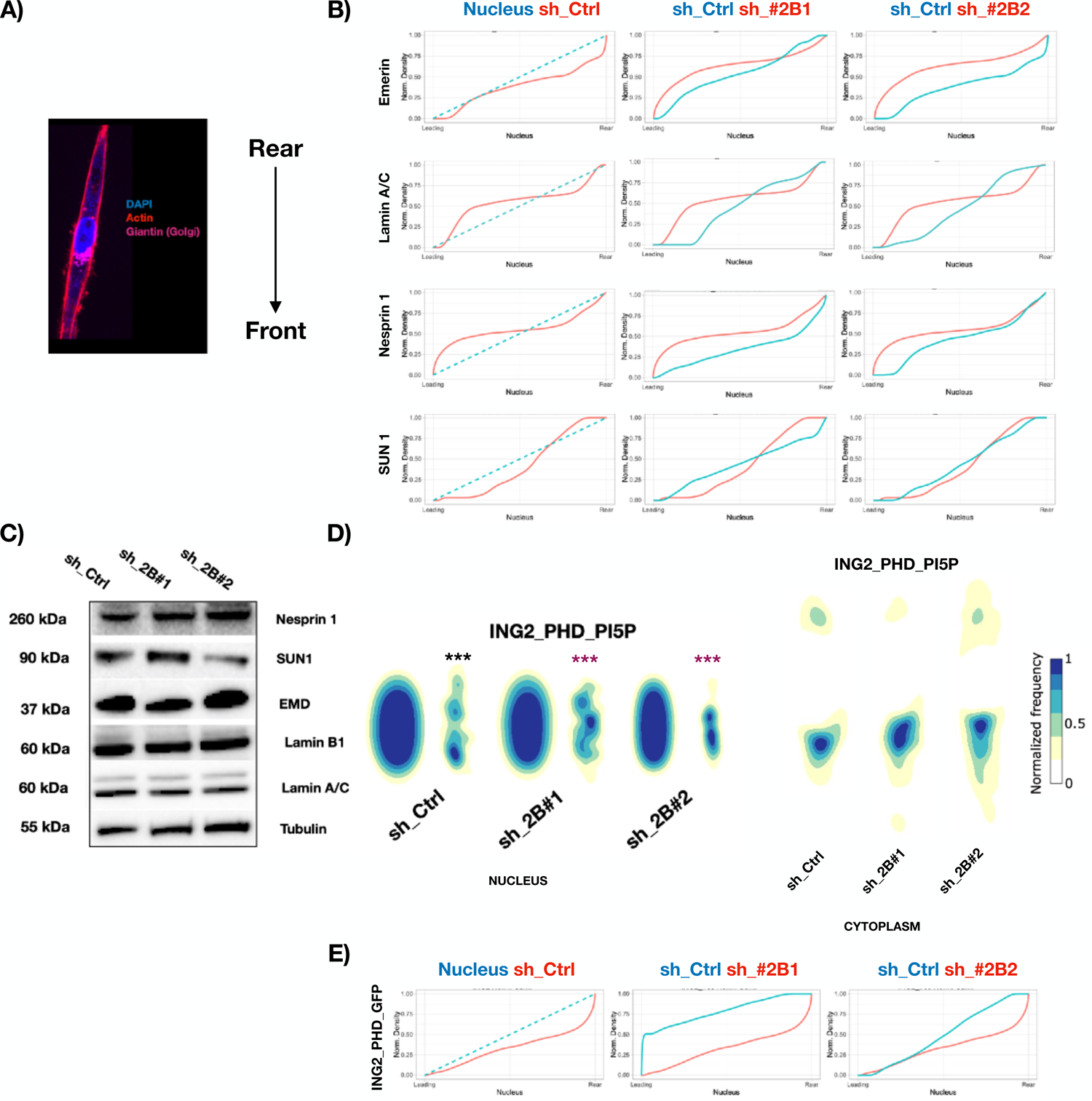
A) Immunofluorescence staining for Actin (Phalloidin), nucleus (DAPI) and Golgi (Giantin) in hTERT_RPE1 seeded on FN-coated coverslips displaying linear (10um width) PLL-g-Peg patterns. Orientation of the cells (Front/Rear) was assessed considering Golgi localisation always at the front. B) Protein distribution plots of normalized density, corresponding to maps in Figure 2F. C) Western Blotting shooing expression of nuclear envelope proteins in hTERT_RPE1 cells transduced to silence PIP4K2B (sh_2B#1/#2), or with empty pLKO_1 vector as control (sh_Ctrl). D) Orientation maps of nuclear (left) and cytoplasmic (right) localisation of PI5P, detected using ING2_PHD-GFP probe. Cells were transduced to silence PIP4K2B (sh_2B#1/#2, n=36/51), or with empty pLKO_1 vector as control (sh_Ctrl, n=50). E) Protein distribution plots of normalized density, corresponding to maps in Supplementary Figure 2D. Statistical analysis was performed using Kolmogorov–Smirnov test for nuclear front/rear enrichment. Black * indicates statistical analysis of the front vs rear accumulation, while purple * indicates statistical analysis of Ctrl cells vs PIP4K2B knock-down cells.

**Supplementary Figure 3.**
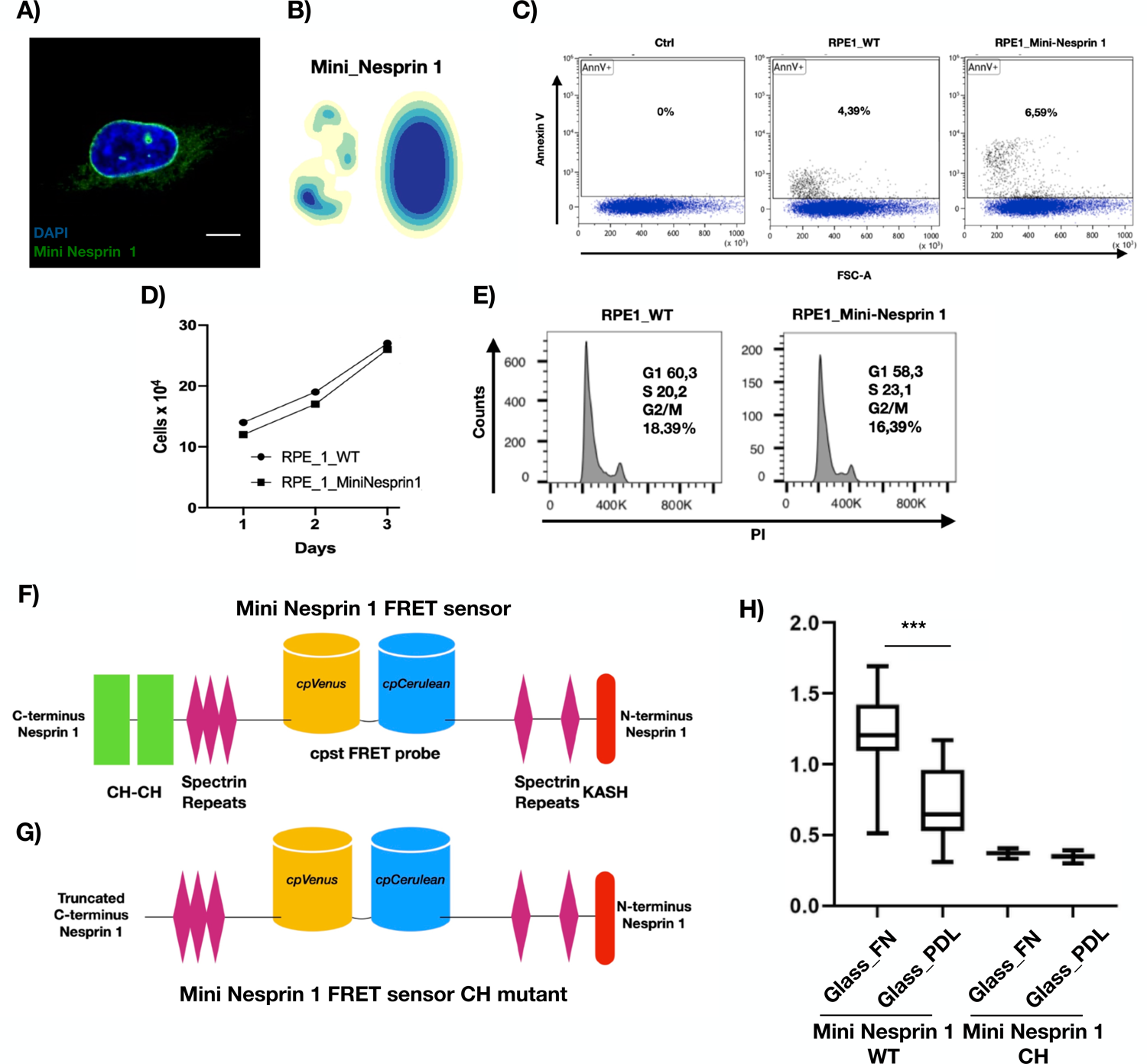
A) Immunofluorescence staining for Mini Neprin 1 cpst-FRET sensor (green) and nucleus (DAPI) (scale bar = 10μm). B) Orientation maps of nuclear localisation of Mini Neprin 1 cpst-FRET sensor. C) FACS based Annexin-V assay to assess apoptosis in hTERT_RPE1 cells wild type (RPE1_WT) or transfected to express Mini Neprin 1 cpst-FRET sensor (RPE_1 Mini_Nesprin 1). D) Cell proliferation assay performed in 24 well plates. 104 hTERT_RPE1 cells wild type (RPE1_WT) or transfected to express Mini Neprin 1 cpst-FRET sensor (RPE_1 Mini_Nesprin 1) were plated and then manually counted for 3 days. E) FACS based cell cycle analysis employing propidium iodide (PI) incorporation in the cells of hTERT_RPE1 cells wild type (RPE1_WT) or transfected to express Mini Neprin 1 cpst-FRET sensor (RPE_1 Mini_Nesprin 1). F)/G) Graphical representation of Mini Neprin 1 cpst-FRET sensor (top) and CH-CH mutant (bottom). H) Nuclear envelope (NE) tension analysis exploiting Mini Nesprin 1 cpst-FRET sensor wild-type version (WT) or CH-CH_mutant (CH). hTERT_RPE1 cells expressing either WT or CH versions of the FRET sensor were seeded on FN-coated glass or soft surfaces (Gel 2.3 kPa). Data quantification is shown as boxplot chart representing inverted FRET index values (donor/acceptor, the higher the value, the higher the tension) and plotted as boxpolots using Tukey’s method in prism7 software. Statistical analyses were performed using unpaired Student’s t test with Welch’s correction, *P < 0.05, **P < 0.01, **P < 0.001.

**Supplementary Figure 4.**
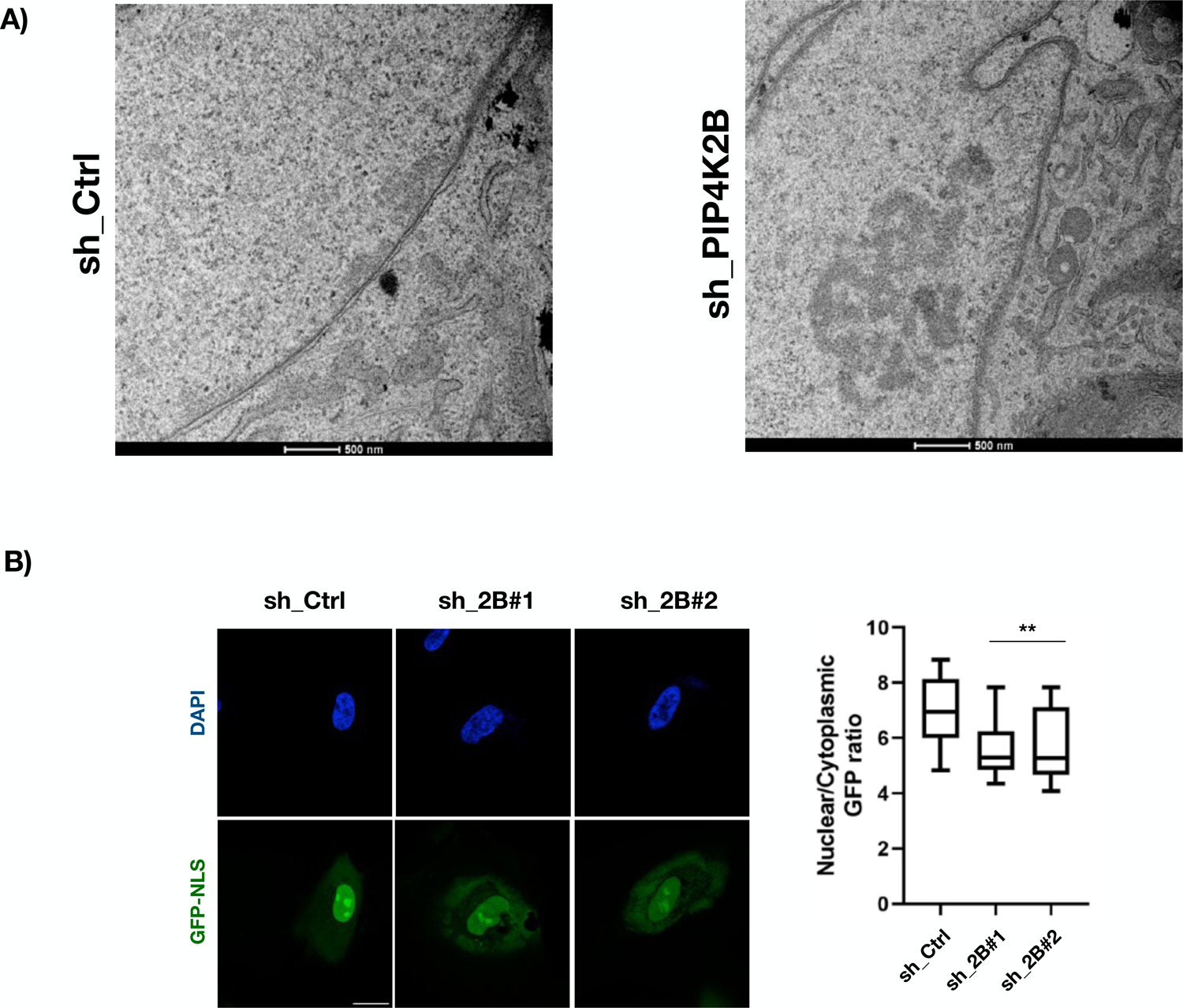
A) TEM images of sh_Ctrl or sh_PIP4K2B cells. Cropped images are presented in Figure 2 (scale bar = 500nm). B) Nuclear import assay. hTERT_RPE1 cells were transduced to silence PIP4K2B (sh_2B#1/#2), or with empty pLKO_1 vector as control (sh_Ctrl). Next, they were transfected with a plasmid encoding a GFP tagged with a single nuclear exportation signal (NLS), which mildly drive GFP localisation into the nuclei (scale bar = 10μm). Nuclear/Cytoplasmic ratio of GFP intensity was analysed and plotted as boxplots using Tukey’s method in prism7 software. Statistical analyses were performed using unpaired Student’s t test with Welch’s correction, *P < 0.05, **P < 0.01, **P < 0.001.

**Supplementary Figure 5.**
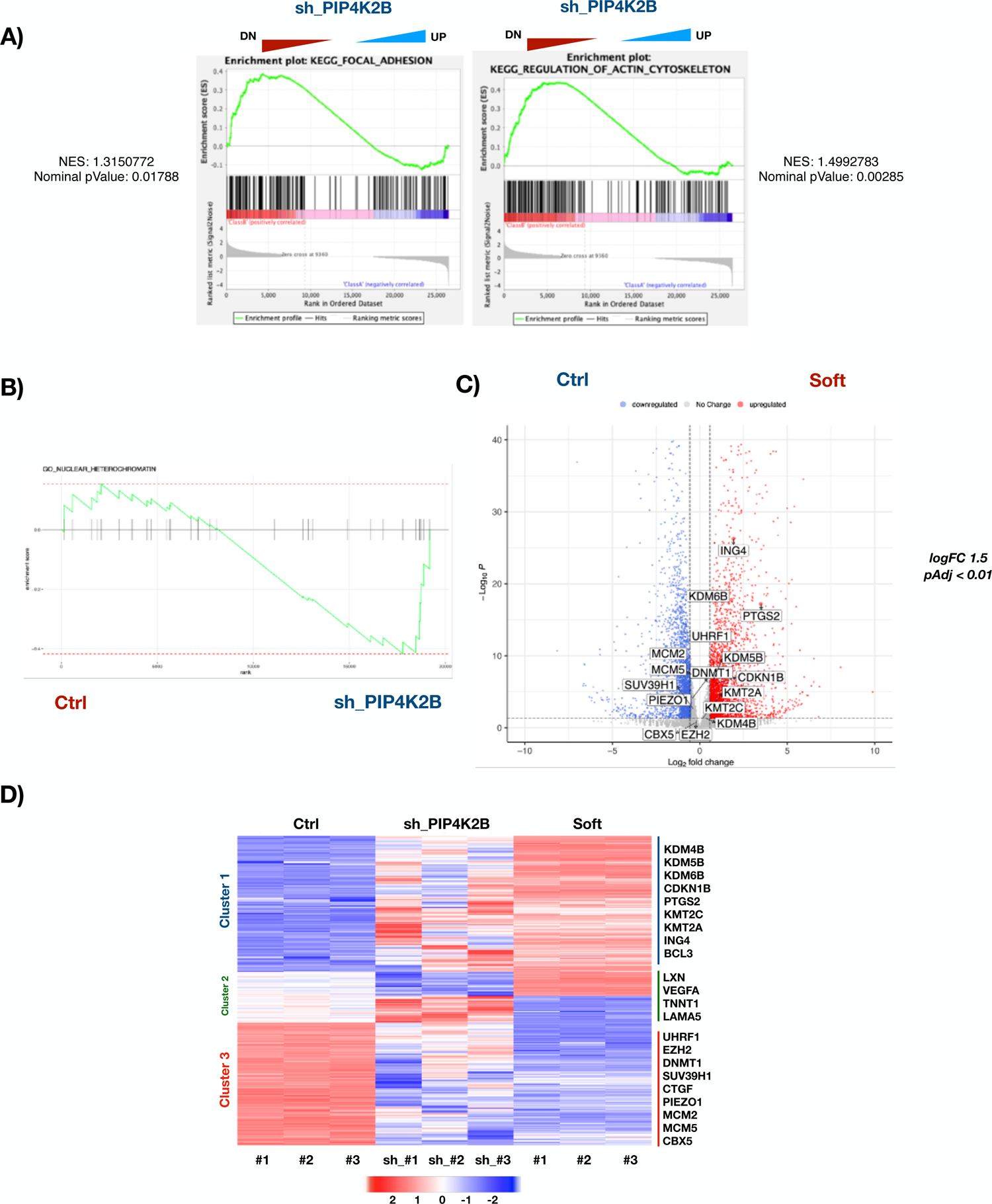
A) Gene expression data generated through RNA-seq comparing Ctrl vs sh_PIP4K2B were used for GSEA analyses to extract biological knowledge. False Discovery Rate (FDR) Normalised Enrichment Score (NES) are reported. B) Enrichment plot of GO nuclear heterochromatin terms. The green curve corresponds to the ES (enrichment score) curve, which is the running sum of the weighted enrichment score obtained from fGSEA software. C) Volcano plot representing DEGenes of Soft vs Ctrl conditions. D) Heatmap of DEGenes found in both sh_PIP4K2B and Soft compared to Ctrl. Genes are presented in 3 clusters: 1) Upregulated genes in both sh_PIP4K2B and Soft; 2) Genes differentially regulated in Soft and sh_PIP4K2B; 3) genes Downregulated in both sh_PIP4K2B and Soft.

**Supplementary Figure 6.**
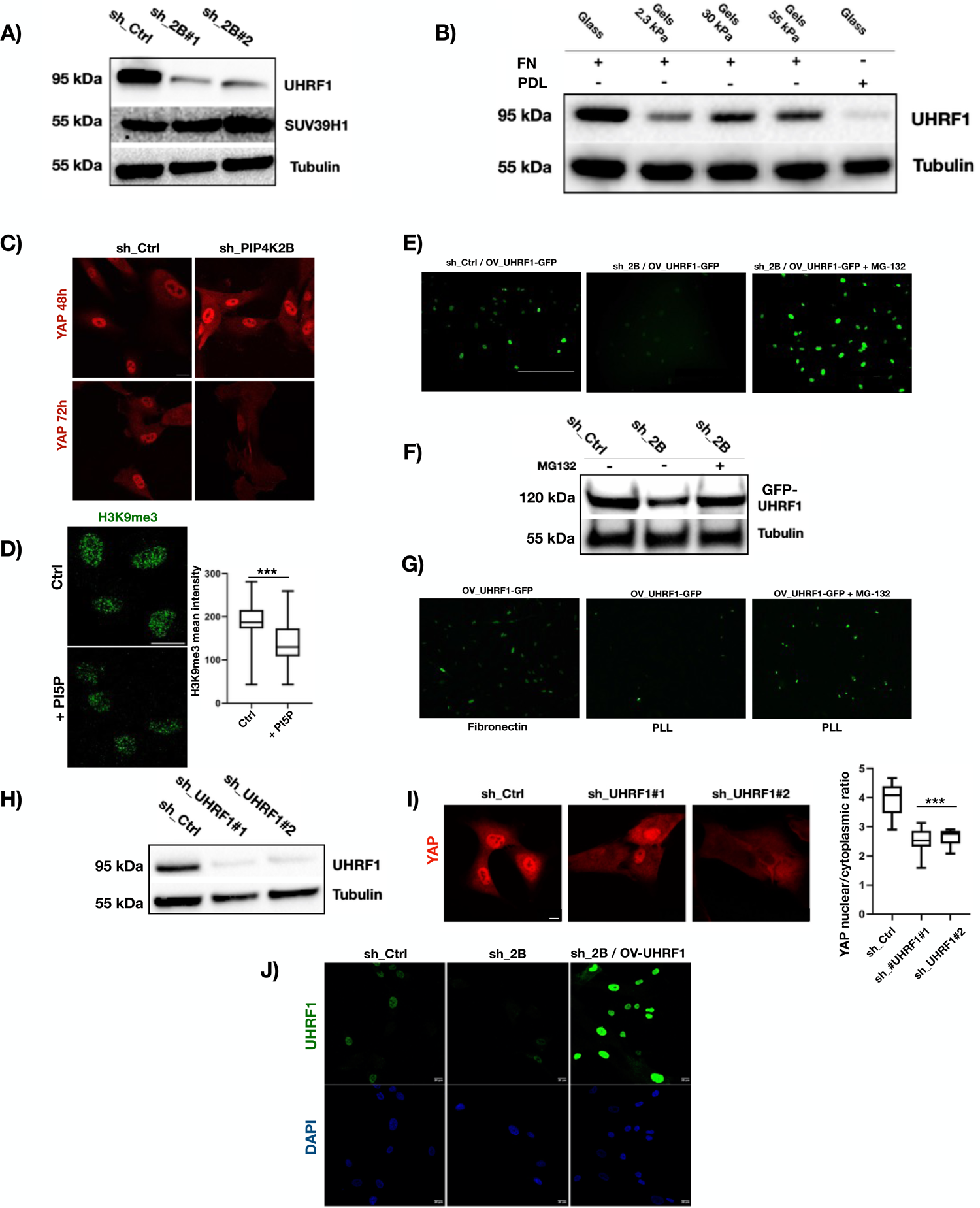
A) Western Blotting of hTERT_RPE1 cells transduced to silence PIP4K2B (sh_2B#1/#2), or with empty pLKO_1 vector as control (sh_Ctrl). Immunoblot was performed to detect protein levels of UHRF1 and SUV39H1. Tubulin was used as loading control. B) hTERT_RPE1 cells were cultured for 24 hours on Fibronectin (FN)- or Poly D-lysine (PDL)-coated glass coverslips, and on FN-coated acrylamide gels of different stiffness (2.3 kPa / 30 kPa / 55 kPa), then lysed. Protein lysates were immunoblotted to analyse UHRF1 protein levels. Tubulin was used as loading control. C) Immunofluorescent staining of YAP cellular distribution in sh_Ctrl and sh_PIP4K2B cells related to WB time course experiment presented in Figure 4A (scale bar = 10μm). Quantification of nuclear to cytoplasmic YAP signal ratio are reported as boxplots plotted using Tukey’s method in prism7 software. D) Immunofluorescence staining of H3K9me3 in nuclei of cells starved overnight and then grown for 4 hours with (+PI5P) or without PtdIns5P (Ctrl) (scale bar = 20μm). Data quantification is shown as boxplots representing the ratio between the intensity of single fluorescent dots and cell nuclear area. E)/F) Immunofluorescence and Western Blotting of hTERT_RPE1 cells transduced to silence PIP4K2B (sh_2B) growing with or without MG-132 (5μM), or with empty pLKO_1 vector as control (sh_Ctrl) (scale bar = 400μm). Cells were previously transduced to overxpress GFP tagged-UHRF1. IF to detect GFP-UHRF1 levels and Immunoblot were performed to detect protein levels of exogenous UHRF1. Tubulin was used as loading control. G) Immunofluorescence analysis of GFP-UHRF1 in cells seeded on Fibronectin or PDL (+/- MG-132, 5 μm) coated glass coverslips. H) Western Blotting of hTERT_RPE1 cells transduced to silence UHRF1 (sh_UHRF1#1/#2) or with empty pLKO_1 vector as control (sh_Ctrl). Protein levels of UHRF1 were assessed, using Tubulin as loading control. I) Immunofluorescent staining of YAP cellular distribution in cells treated as in G) (scale bar = 10μm). Quantification of nuclear to cytoplasmic YAP signal ratio are reported as boxplots plotted using Tukey’s method in prism7 software. J) Immunofluorescent staining of UHRF1 in cells transduced with empty pLKO_1 vector (sh_Ctrl), with sh_RNAi targeting PIP4K2B (sh_PIP4K2B), and with sh_RNAi targeting PIP4K2B and a plasmid to overexpress UHRF1 (sh_PIP4K2B / OV_UHRF1) (scale bar = 20μm).

**Supplementary Figure 7.**
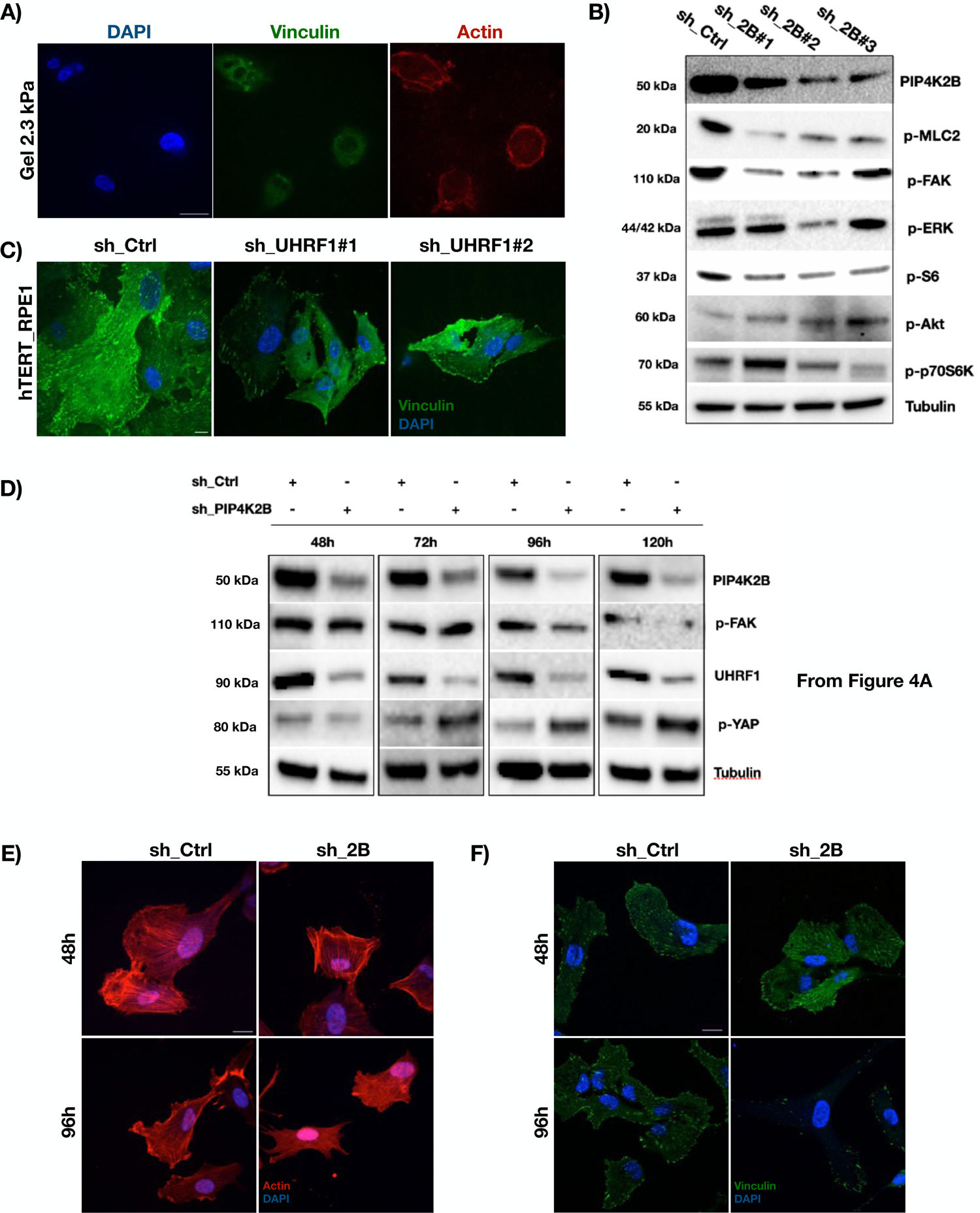
A) Immunofluorescent staining of nuclei (DAPI), Focal Adhesion (Vinculin) and Actin (Phalloidin) in hTERT_RPE1 cells seeded on FN-coated soft surfaces (Gel 2.3 kPa) (scale bar = 20μm). B) Western Blotting analysis of protein levels of protein and phosphorylated proteins in cells depleted for PIP4K2B (sh_2B#1/#2/#3) and control cells (sh_Ctrl). Tubulin was used as loading control. C) Immunofluorescence staining of focal adhesions (Vinculin) in in hTERT_RPE1 cells seeded on FN-coated glass and transduced to silence UHRF1 (sh_UHRF1#1/#2) or with empty pLKO_1 vector as control (sh_Ctrl) (scale bar = 10μm). D) Western Blotting analysis of phosphorylated FAK levels. WB is related to WB shown in Figure 4A. E)/F) Immunofluorescence staining of Actin (Phalloidin) or focal adhesions (Vinculin) in cells treated as in D). Data are related to WB shown in D) (scale bar = 10μm).

**Supplementary Figure 8.**
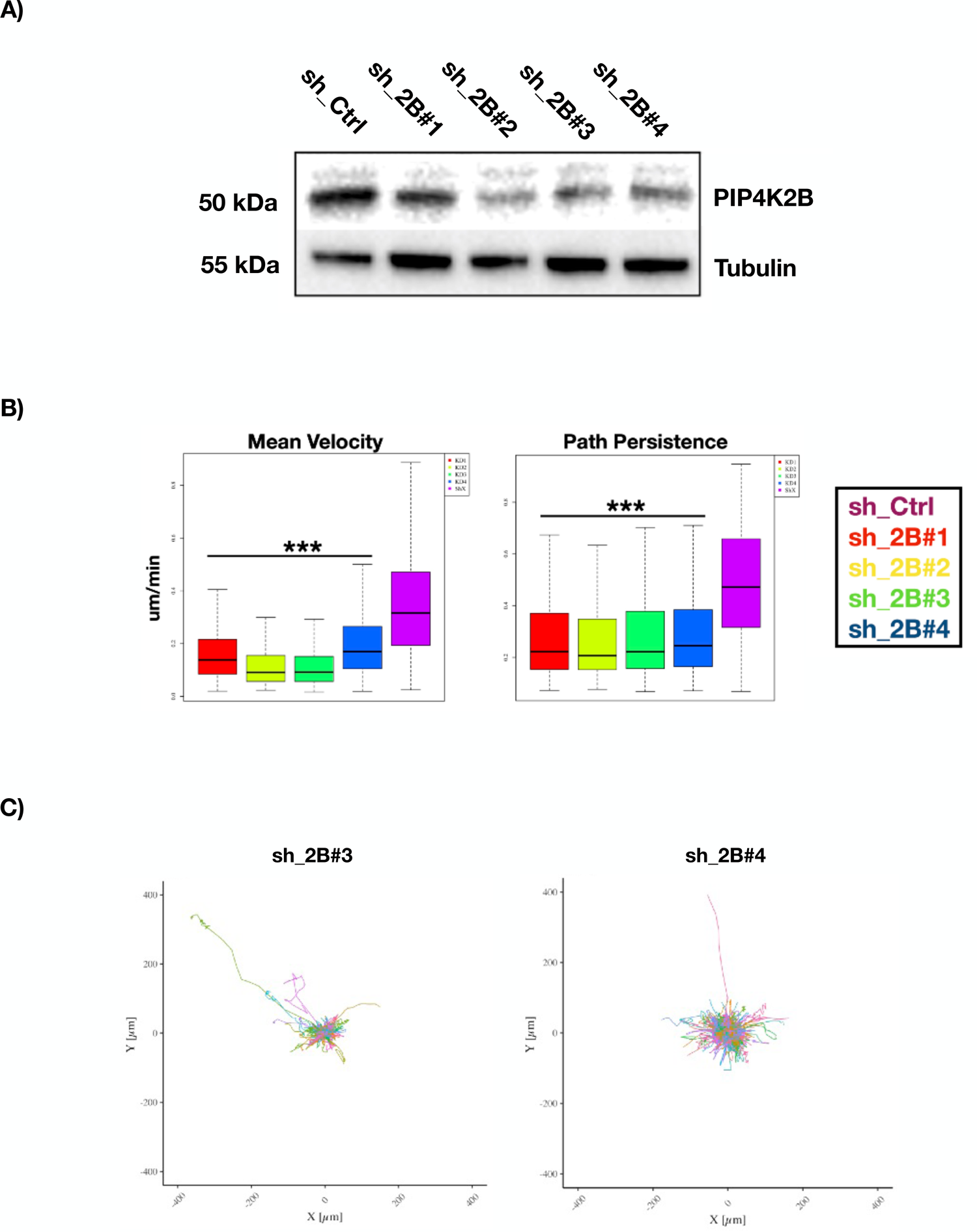
A) Western Blotting analysis of PIP4K2B protein expression in cells transduced with 4 different sh_RNAi targeting PIP4K2B (sh_2B#1/#2/#3/#4). Empty pLKO_1 vector was used as control (sh_Ctrl) and Tubulin as loading control. B) 2D cell motility quantification of cell Mean Velocity and Path Persistence. Cells were treated as in A), and seeded on FN-coated glass coverslips. Nuclei were stained with NucBlue and cell tracking performed for 16 hours.

## Notes

### Competing Interest Statement

The authors have declared no competing interest.

